# Melatonin modulates habituation learning via the convergent action of two MT_1_-type receptors

**DOI:** 10.64898/2026.09.10.749335

**Authors:** Dominique Baas, Isabelle Darvaux-Hubert, Romane Dorado-Doncel, Andrew Hsiao, Lisa Broisin, Abdel Rahman El Hassan, Julien Perrichet, Owen Randlett

**Affiliations:** Laboratoire MeLiS, Université Claude Bernard Lyon 1 - CNRS UMR5284 - Inserm U1314, Faculté de Médecine et de Pharmacie, 8 avenue Rockefeller, 69008 Lyon, France; Master program: Bioscience, Ecole Normale Supérieure de Lyon, France; Licence program: Science de la Vie, Université Claude Bernard Lyon 1, Lyon, France

## Abstract

Melatonin is a hormone produced by the pineal gland and retina, with roles in sleep and circadian rhythms as well as other neuronal processes, including the regulation of learning and memory. Using larval zebrafish, we previously identified Melatonin as a potent modulator of habituation learning: the progressive suppression of responses to a repeated stimulus. Here we have extended these analyses and have found that Melatonin has strong effects on separate aspects of habituation learning: it potentiates habituation of response probability and latency, while simultaneously inhibiting habituation of movement amplitude and duration. We performed a systematic CRISPR mutagenesis of all six zebrafish Melatonin receptors, and found that only two MT_1_-type receptors (Mtnr1aa and Mtnr1al) are required for Melatonin’s effects on habituation. These receptors showed a cooperative interaction, with each individual mutant showing reduced sensitivity to Melatonin, and *mtnr1aa;mtnr1al* double mutants showing complete insensitivity. To map where these receptors act, we used whole-brain activity mapping. Despite distinct footprints, both perturbed a shared set of brain areas, including the torus longitudinalis, cerebellum, and preoptic area, with the double mutant producing the most extensive phenotype by combining and expanding the effects seen in each single mutant. Thus, we propose that two MT_1_-type receptors act at overlapping circuit nodes to reinforce Melatonin’s bidirectional control of learning.

## Introduction

Habituation is an evolutionarily conserved form of learning, defined as a progressive decrease in responsiveness to a repeatedly presented stimulus (Rankin et al., 2009). Habituation displays a well-defined set of parametric properties: it is stimulus-specific, recovers spontaneously after a rest period, is accelerated by high-frequency stimulus presentation, and can be disrupted by the introduction of a novel stimulus (Thompson and Spencer, 1966). Far from being a passive consequence of neural fatigue or sensory adaptation, habituation is an active form of neural plasticity, involving persistent changes in multiple systems (Randlett et al., 2019; McDiarmid et al., 2019; Ramaswami, 2014; Cooke and Ramaswami, 2020; Nelson et al., 2023). Deficits in habituation are associated with neurodevelopmental and psychiatric conditions including autism spectrum disorder, schizophrenia, and ADHD (McDiarmid et al., 2017), underscoring its relevance to normal cognitive function and its potential as a tractable behavioural readout of circuit-level pathology.

Larval zebrafish have emerged as a powerful system for dissecting the neural and molecular substrates of habituation learning. When exposed to a sudden transition to darkness (a dark flash, DF), larvae execute a stereotyped escape maneuver called the O-bend (Burgess and Granato, 2007), which habituates robustly with repeated stimulation (Wolman et al., 2011; Randlett et al., 2019). The transparency and genetic tractability of larvae, combined with whole-brain activity mapping and calcium imaging approaches, have enabled identification of distributed plasticity across multiple brain regions that collectively drive DF habituation learning (Randlett et al., 2019; Lamiré et al., 2023). Furthermore, the accessibility of larval zebrafish to high-throughput pharmacological and genetic screening has allowed systematic identification of neuromodulatory pathways that regulate habituation performance (Wolman et al., 2011, 2015). Through such a screen, we previously identified Melatonin as one of the most potent modulators of habituation learning, with acute exposure causing a dramatic enhancement of habituation for the probability of executing an O-bend response to DFs (Lamiré et al., 2023).

Melatonin is a neuroendocrine hormone synthesized from tryptophan via serotonin, with the rate-limiting step catalyzed by arylalkylamine N-acetyltransferase (AANAT) (Ganguly et al., 2002). Its production is under tight circadian control, resulting in high nocturnal concentrations in the blood and cerebrospinal fluid across vertebrate species, including zebrafish (Falcón et al., 2009; Cahill, 1996; Gandhi et al., 2015). In zebrafish, two AANAT paralogs contribute to Melatonin synthesis: Aanat1, expressed primarily in the retina, and Aanat2, expressed in both the retina and the pineal gland (Gandhi et al., 2015; Hill et al., 2026). This positions Melatonin as a key hormonal signal coupling the external light/dark cycle to internal physiological state.

Melatonin acts primarily through two G-protein coupled receptors in the mammalian brain, MT_1_ (Mtnr1a) and MT_2_ (Mtnr1b), which signal through Gi proteins to reduce intracellular cAMP, though they exhibit distinct binding kinetics, signaling properties, and expression patterns (Dubocovich and Markowska, 2005; Liu et al., 2016; Cecon et al., 2017). Both receptors are broadly expressed in the brain, including regions implicated in circadian regulation, sleep, and plasticity (Ng et al., 2017; Klosen et al., 2019). Zebrafish possess an expanded repertoire of six Melatonin receptors: three MT_1_-type (Mtnr1aa, Mtnr1ab, Mtnr1al), two MT_2_-type (Mtnr1ba, Mtnr1bb), and Mtnr1c, an ortholog of the mammalian Melatonin-insensitive GPR50 (Hill et al., 2026). This expanded repertoire provides a potentially powerful genetic handle for dissecting the differential functional effects of Melatonin receptors *in vivo*. A recent study exploiting this genetic handle found that MT_1_-type paralogs mediate Melatonin’s sleep-promoting effects in zebrafish by suppressing behavioural responses to visual stimuli, with Mtnr1aa accounting for most of this effect (Hill et al., 2026).

In addition to its endogenous role, Melatonin is one of the most widely consumed over-the-counter supplements, and its use has increased markedly over the past two decades (Li et al., 2022). In the United States, annual sales nearly tripled between 2016 and 2020, and by 2020 Melatonin had become the substance most frequently ingested by children reported to poison-control centres, with paediatric ingestions rising by 530% over the preceding decade (Lelak et al., 2022). Large numbers of people are therefore exposed to exogenous, often supraphysiological, Melatonin, making it important to understand its actions beyond the promotion of sleep.

Beyond its canonical roles in sleep and circadian entrainment, Melatonin has been increasingly implicated in other cognitive functions, including learning and memory. In rodents, MT_1_/MT_2_ signaling shapes time-of-day-dependent hippocampal synaptic plasticity and learning efficiency (Jilg et al., 2019). Individual receptor knockouts reveal distinct functional roles, consistent with the largely non-overlapping neuronal expression of MT_1_ and MT_2_ in the brain (Klosen et al., 2019): MT_2_-deficient mice show impaired hippocampal long-term potentiation (Larson et al., 2006), and loss of MT_2_ precludes the memory-enhancing effect of chronic Melatonin treatment on long-term recognition memory, while loss of MT_1_ only partially mitigates this effect (Pistono et al., 2021). Additionally, deletion of either MT_1_ or MT_2_ alone is sufficient to abrogate reward-associated conditioned place preference (Clough et al., 2014). However, genetic deletion of both receptors has been reported to *enhance* cognitive and motor performance (O’Neal-Moffitt et al., 2014), in apparent contradiction with the impairment phenotypes described above. This suggests that the consequences of MT_1_/MT_2_ signaling in rodents are strongly paradigm- and context-dependent, rather than reflecting a single, generalizable role in cognition. Therefore, Melatonin receptor signaling appears to have important roles in plasticity that are separable from its role in sleep promotion, but the circuit mechanisms and specific receptor subtypes responsible have not been well-defined.

Here we leverage the zebrafish system to address this gap. Using a large behavioural dataset, we first show that Melatonin has complex, bidirectional effects on multiple independently regulated components of habituation behaviour. We then demonstrate through systematic CRISPR mutagenesis of all six zebrafish Melatonin receptors that two MT_1_-type receptors, Mtnr1aa and Mtnr1al, cooperatively and non-redundantly mediate these effects. Finally, using whole-brain activity mapping, we find that although each receptor has a distinct functional footprint, both converge on a shared set of circuit nodes, providing a candidate anatomical basis for their non-redundant, cooperative action. This contrasts with their role in sleep, where Mtnr1aa predominates and Mtnr1ab and Mtnr1al instead act redundantly, partially compensating for its loss (Hill et al., 2026), indicating that the same small family of receptors can be deployed with contrasting genetic logic. Together, these data provide a molecular and functional dissection of how Melatonin modulates a fundamental form of learning through the convergent action of receptor paralogs.

## Results

### Melatonin induces bidirectional changes in visual habituation learning

We previously identified Melatonin as a potent modulator of habituation learning in larval zebrafish (Lamiré et al., 2023). Based on this previous dataset, we concluded that Melatonin potently enhances habituation learning for the probability of executing an O-bend response to repeated DF stimuli. Importantly, we found that Melatonin had little effect on the habituation of O-bend displacement, which is governed by a distinct plasticity mechanism (Randlett et al., 2019), and that Melatonin did not affect the response of the larvae to acoustic or optomotor stimuli (Lamiré et al., 2023). This demonstrates that Melatonin does not have a generalized effect on sensory processing or motor output (e.g. by simply inducing sleep Gandhi et al., 2015; Hill et al., 2026).

Here, we have extended these analyses using a larger dataset, and confirmed that acute exposure to 1 µM Melatonin potently enhances habituation learning for the probability of response (**Figure 1A**). The response to the first 3 dark-flash stimuli are not dramatically altered by the treatment (“Naive Response”: ***Figure S1Aii***), but the effect is pronounced during the remaining 57 flashes of the first training block (“Block 1” ***Figure S1Aiii***), the 180 flashes in the subsequent 3 training blocks (“Trained Response”: ***Figure S1Aiv***), and the re-test block 5 hours after the last training session (“Block 5”: ***Figure S1Avi***). Consistent with our previous data, the response of the larvae to the acoustic stimuli was indistinguishable from controls (“Acoustic Response”: **Figure 1Ai**,***Figure S1Av***; Lamiré et al., 2023), confirming that the effect of Melatonin is not one of generalized sedation. From these data we can conclude that Melatonin potentiates habituation learning for the probability of response.

**Figure 1.**
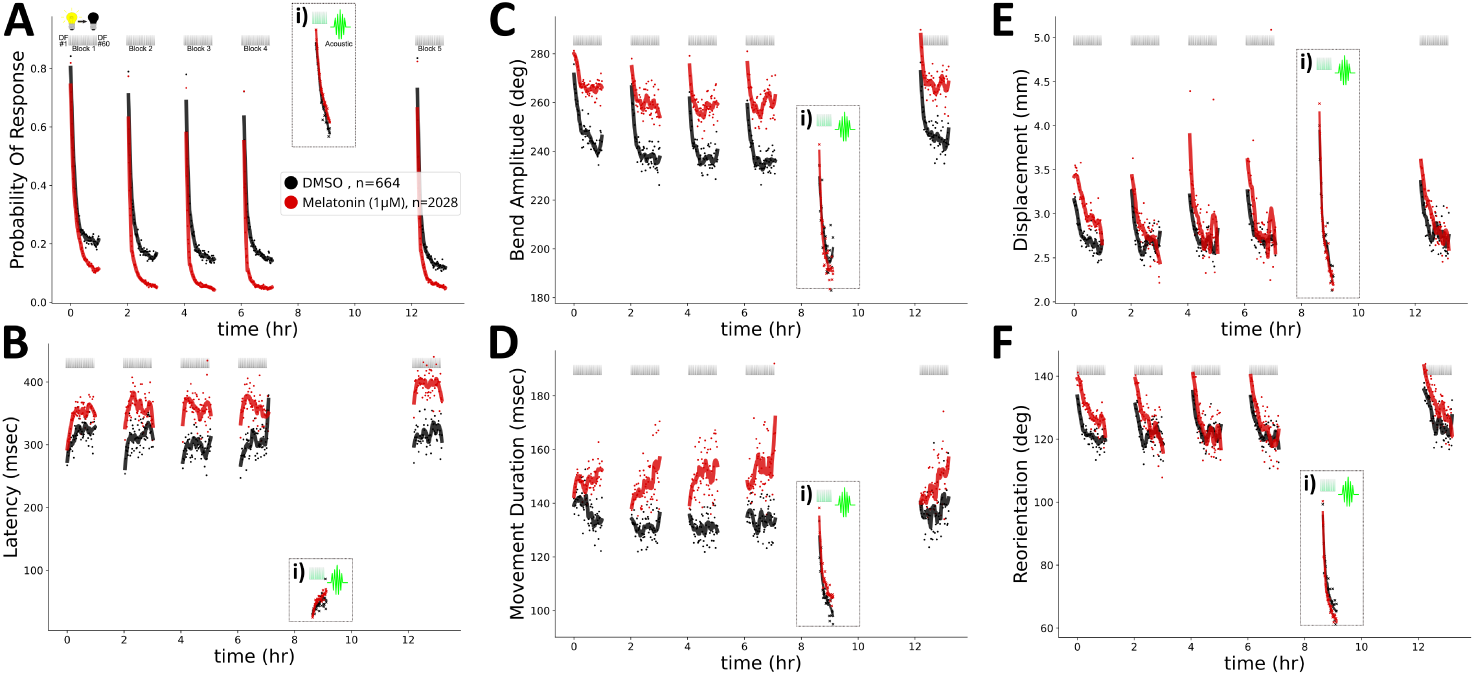
Melatonin has bidirectional effects on different components of habituation learning A) Treatment with 1 µM Melatonin (red) results in more rapid and profound decreases in the probability of response to DF stimuli during habituation training relative to vehicle controls (0.1% DMSO black; n = 664 vehicle and n = 2188 Melatonin larvae, 18 different plates). DF stimuli are delivered at 1-minute intervals, in 4 blocks of 60 stimuli, separated by 1 hour of rest (from 0:00-7:00). 1.5 hours later a block of 30 vibration stimuli are delivered at 1-minute intervals (i). Each dot is the mean response across larvae to each DF, while ‘X’s mark mean responses to vibration stimuli. Lines are smoothed in time with a Savitzky–Golay filter (window = 15 stimuli, order = 2). B-F) Same as A, but for the kinematic components of the response: B) the response latency, C) the maximum tail-bend amplitude achieved during the movement, D) the duration of the movement, E) the displacement achieved, and F) the reorientation achieved by the larvae.

This larger dataset now also allowed us to more carefully examine the effects of Melatonin on the behavioural components related to the magnitude of the O-bend response. Indeed, a significant enhancement of habituation was also observed for the response latency, which increases by 50-100 msec over habituation training (**Figure 1B, *Figure S1B***). This effect was also visible in our original datasets (Lamiré et al., 2023), but was not previously clear and significant, likely due to the higher degree of variability in these kinematic behavioural measures relative to the probability component, as well as the fact that the strong suppression of responses by Melatonin results in little data for the kinematic measures, as these can only be observed when fish respond to the stimuli.

Interestingly, while the probability and latency components of habituation learning were enhanced by Melatonin, the opposite was observed for bend amplitude (the maximum tail-angle bending achieved during the O-bend movement) (**Figure 1C, *Figure S1C***), as well as the movement duration (the time taken from the initiation of the movement until its termination) (**Figure 1D, *Figure S1D***). Both of these components normally decrease with training, resulting in movements with reduced amplitudes. This effect was significantly inhibited by Melatonin, where treated-larvae maintain more vigorous movements across stimulus presentations. As with probability and latency, the naive response is much less affected than the response seen during training, indicating that the primary effect is on habituation of the response, rather than their innate sensitivity.

Finally, while Melatonin inhibits habituation of bend amplitude and movement duration, we did not observe such clear alterations when analyzing the habituation performance curves for displacement (the displacement of the larvae during the O-bend movement) (**Figure 1E, *Figure S1E***) or reorientation (the change in heading direction achieved by the larvae during the O-bend movement) (**Figure 1F, *Figure S1F***), consistent with our previous finding that Melatonin has little effect on displacement habituation (Lamiré et al., 2023). Treated- and untreated-larvae showed similar modulation of these parameters, resulting in little divergence in their performance curves across habituation training. However, in these two components, the response is subtly but consistently increased in the Melatonin-treated group, primarily in the “Naive” and “Block 1” response, indicating that Melatonin has a weak stimulatory effect on these aspects of the movements, but does not appear to have a substantial effect on the habituation process.

Collectively, these behavioural analyses reveal that Melatonin has a complex and bidirectional effect on different components of habituation learning, potentiating some aspects of learning, while inhibiting others, and leaving others unaffected. This is consistent with a model in which Melatonin acts on specific neural circuits that govern distinct components of habituation learning, and cannot be explained by a generalized effect, like modulating sleep/arousal state, or visual sensitivity.

### Zebrafish larvae lacking Melatonin production show habituation deficits for the probability of response

Our pharmacological experiments have shown that exogenously applied Melatonin has a potent effect on habituation learning, but the relevance of endogenous Melatonin is not clear. To address this question, we tested zebrafish mutants that are unable to produce Melatonin due to mutations in Arylalkylamine N-acetyltransferase (Aanat). Aanat is encoded by two genes in zebrafish, *aanat1* and *aanat2*, which are expressed in the retina, and the retina and pineal gland, respectively (Gandhi et al., 2015; Hill et al., 2026). If endogenously produced Melatonin influences habituation in the same ways as exogenous Melatonin, we would expect these Melatonin-deficient mutants to exhibit learning deficits for probability and latency, and increased learning performance for bend amplitude and movement duration. Consistent with this hypothesis, we found that double mutants (*aanat1^-/-^*; *aanat2^-/-^*) show a significant impairment in habituation learning for the probability of response (**Figure 2A, *Figure S2A***). Similarly to the effect of exogenous Melatonin, this effect is most pronounced during the later stages of training, and is not significant for the naive response to the first 3 dark-flash stimuli. Notably, this effect was also only observed in the double mutants. This redundancy demonstrates either that both Aanat1 and Aanat2 contribute to the production of Melatonin that promotes habituation learning, or that there is functional redundancy/compensation between these genes.

**Figure 2.**
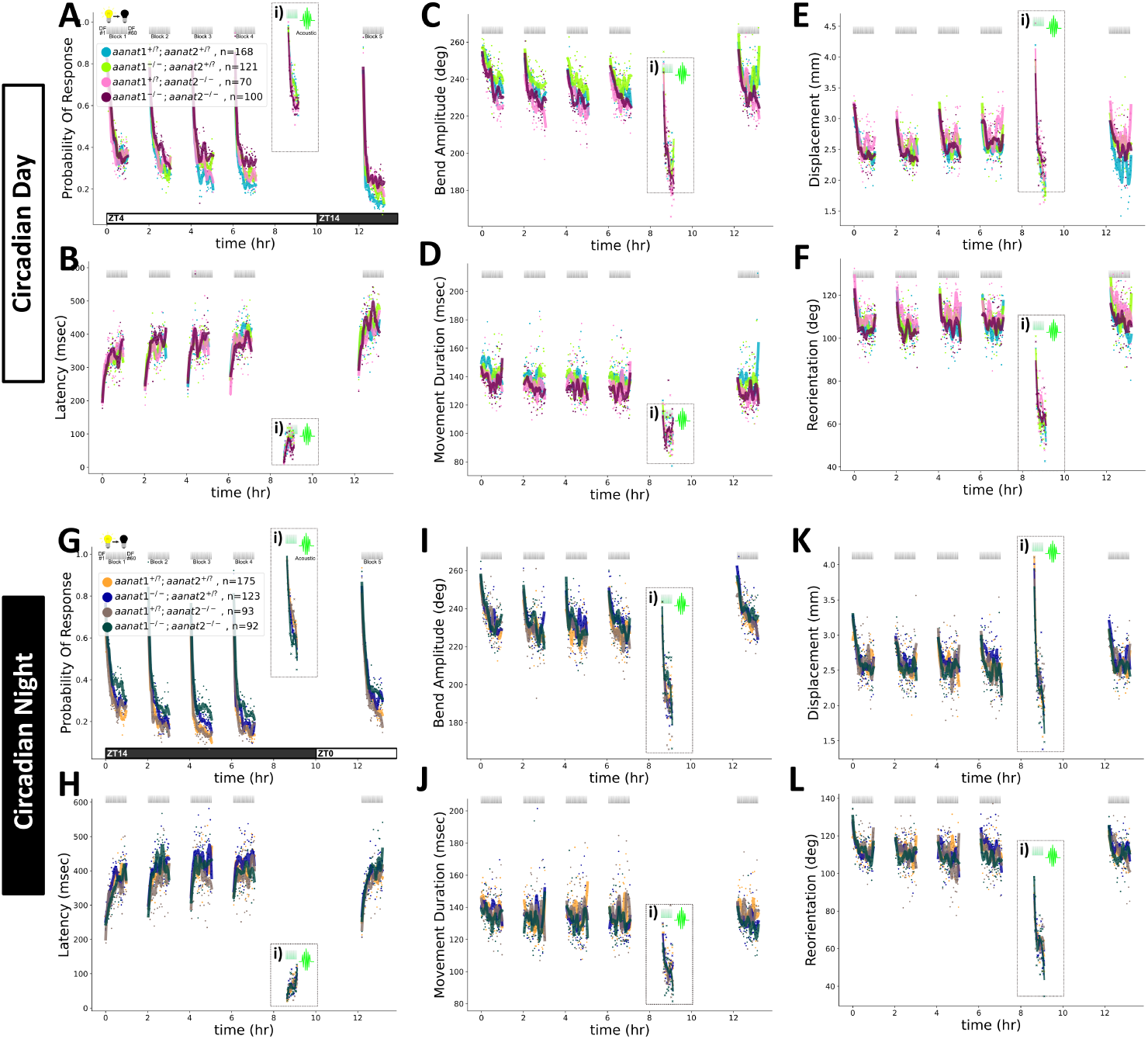
Zebrafish larvae lacking Melatonin production show habituation deficits for the probability of response A) Double mutants for the Melatonin-synthesizing enzymes Aanat1 and Aanat2 show a decreased modulation of the probability of response to DF stimuli during habituation training relative to heterozygous and WT sibling controls (day-tested cohort, 4 different plates: *aanat1^+/?^;aanat2^+/?^*, n = 168; *aanat1^-/-^; aanat2^+/?^*, n = 121; *aanat1^+/?^;aanat2^-/-^*, n = 70; *aanat1^-/-^; aanat2^-/-^*, n = 100). Testing was performed during the circadian day, beginning at ZT4, where ZT0 is the time of lights “on” and ZT12 is the time of lights “off”. DF stimuli are delivered at 1-minute intervals, in 4 blocks of 60 stimuli, separated by 1 hour of rest (from 0:00-7:00). 1.5 hours later a block of 30 vibration stimuli are delivered at 1-minute intervals (i). Each dot is the mean response across larvae to each DF, while ‘X’s mark mean responses to vibration stimuli. Lines are smoothed in time with a Savitzky–Golay filter (window = 15 stimuli, order = 2). B-F) Same as A, but for the kinematic components of the response: B) the response latency, C) the maximum tail-bend amplitude achieved during the movement, D) the duration of the movement, E) the displacement of the movement, and F) the reorientation achieved by the larvae. G) Double mutants for the Melatonin-synthesizing enzymes Aanat1 and Aanat2 show a decreased modulation of the probability of response to DF stimuli during habituation training relative to heterozygous and WT sibling controls (night-tested cohort, shifted light cycle, 4 different plates: *aanat1^+/?^;aanat2^+/?^*, n = 175; *aanat1^-/-^; aanat2^+/?^*, n = 123; *aanat1^+/?^;aanat2^-/-^*, n = 93; *aanat1^-/-^; aanat2^-/-^*, n = 92). H-L) Same as G, but for the kinematic components of the response: H) the response latency, I) the maximum tail-bend amplitude achieved during the movement, J) the duration of the movement, K) the displacement of the movement, and L) the reorientation achieved by the larvae.

Contrary to our expectations, habituation-dependent modulation of the latency component of the O-bend response, as well as the other kinematic parameters, were not clearly affected in the *aanat1;aanat2* mutants (**Figure 2B-F, *Figure S2B-F***). However, while the effect size was small, the bend amplitude and movement duration parameters did show a significantly reduced response in double mutants in some epochs (***Figure S2C,D***), consistent with a role for Melatonin in inhibiting this aspect of learning.

These experiments demonstrate that endogenous Melatonin production does influence habituation, but this effect is only convincingly observed for the probability of response. However, these experiments were performed beginning in the morning, when Melatonin production is naturally low, and therefore the effect of endogenous Melatonin on learning may be limited. Also, Melatonin synthesis is suppressed by light. Since our stimulus paradigm requires the animals to be light-adapted to respond to the DF stimulus (Burgess and Granato, 2007), endogenous Melatonin production is expected to be low during our experiments. To try and overcome these limitations, we also performed experiments in which the larvae were raised in a shifted light cycle, such that they were tested during the circadian night when Melatonin production is higher. However, we observed a similar pattern of results, with the only robust effect again observed for *aanat1;aanat2* double mutants, and only for the probability of response (**Figure 2G-L, *Figure S2G-L***). No significant effect was seen for latency or the kinematic parameters modulated by exogenous Melatonin.

From these data we conclude that endogenous Melatonin does influence habituation learning for the probability of response, but that its role in regulating the other learning parameters is unclear. This may be due to the light-dependent requirements of our visual assay, or it may be that the effects of Melatonin on these other components of habituation learning are only observed when exogenous Melatonin is applied, achieving supra-normal levels of the hormone.

### The effects of Melatonin on habituation learning are mediated by MT_1_-type Melatonin Receptors

As introduced above, zebrafish possess six Melatonin receptors, in contrast to the two (MT_1_ and MT_2_) found in mammals (Hill et al., 2026). To identify which of these receptors mediate the effects of Melatonin on habituation learning, we generated CRISPR mutants for each (***Table 1***), and tested their effect on the modulation of habituation learning by exogenous Melatonin. Since the endogenous effects of Melatonin are subtle (**Figure 2**), we focused our analyses of mutants on the effects of exogenous Melatonin, reasoning that mutants for the relevant receptor(s) should be insensitive to Melatonin treatment.

**Table 1.**
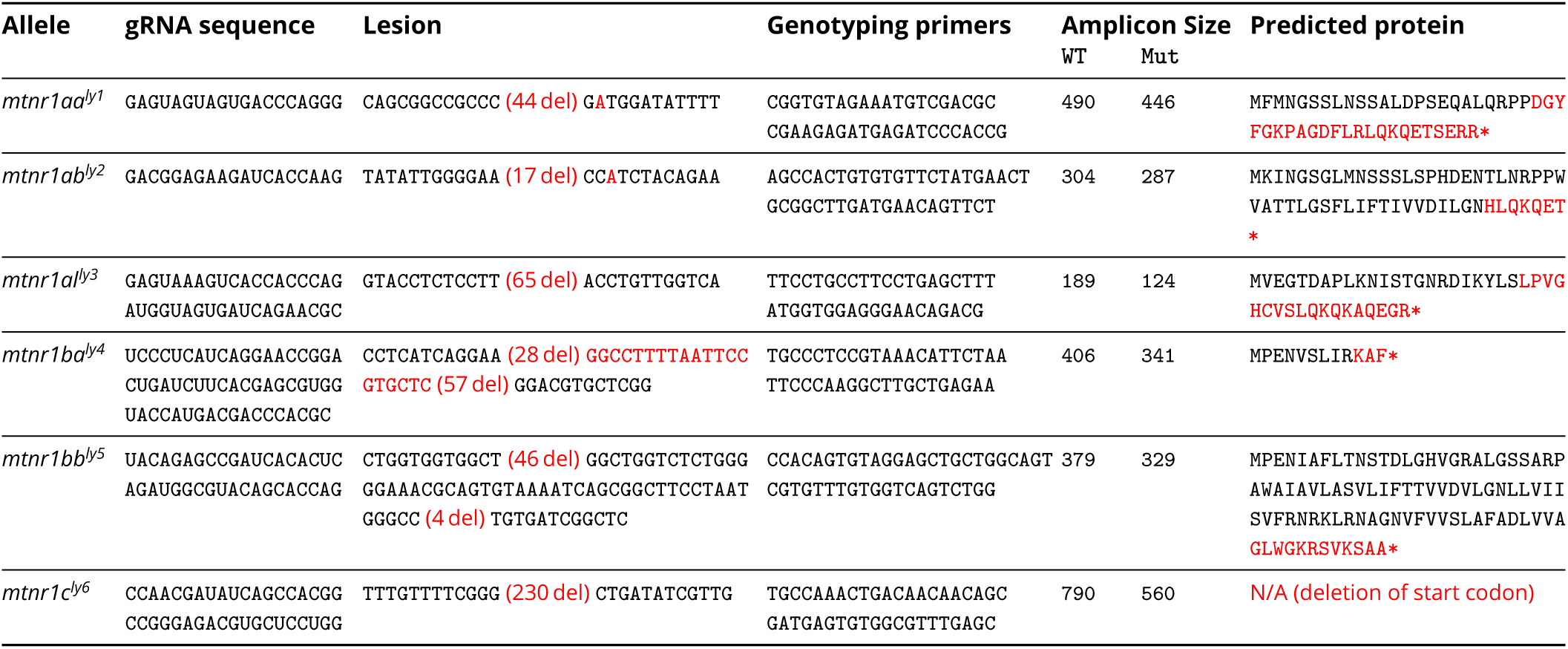
Genotyping primers and allele information for CRISPR mutant lines. Primer sequences and expected amplicon sizes (WT and mutant) are given for each allele used in this study. Sequence shows the genomic sequence flanking the deletion or insertion, with the number of deleted or inserted bases in parentheses. Predicted protein shows the translated sequence downstream of the frameshift, with the premature stop codon marked by an asterisk; for *mtnr1c^ly6^* the deletion removes the start codon.

#### Mtnr1aa and Mtnr1al are each partially required for the effects of Melatonin on habituation learning

Mtnr1aa has been reported as the most broadly expressed Melatonin receptor in the larval zebrafish brain (Hill et al., 2026), making it the top candidate for mediating the behavioural effects of Melatonin. Consistent with this hypothesis, we found that *mtnr1aa* mutants are less sensitive to Melatonin than their WT and heterozygous sibling controls (**Figure 3A-F**,***Figure S3A-F***). This was clearest for the probability of response, where Melatonin-treated mutants habituated less strongly than Melatonin-treated sibling controls (**Figure 3A**). However, they still showed a sensitivity to the effects of Melatonin relative to vehicle controls (***Figure S3A***), indicating that Mtnr1aa does not fully mediate the effects of Melatonin on habituation learning. The mutant-dependent effects were less clear for latency, but qualitatively the Melatonin-treated mutants showed less modulation of latency than sibling-treated controls on the response curves (**Figure 3B**), but this did not reach statistical significance (***Figure S3B***). Finally, *mtnr1aa* mutants were also partially resistant to the effects of Melatonin on the bend amplitude and movement duration kinematic parameters, with Melatonin-treated mutants showing a partial effect and a response magnitude intermediate between untreated- and Melatonin-treated-controls during habituation learning (**Figure 3C-D, *Figure S3C-D***). Therefore, while the effects are most clear for the probability component, the trends are consistent across all behavioural components, indicating that this same receptor mediates the effects of Melatonin on all aspects of habituation learning that we have identified. However, it is also clear that these mutants maintain a significant level of Melatonin-responsiveness, indicating that another receptor likely also contributes.

**Figure 3.**
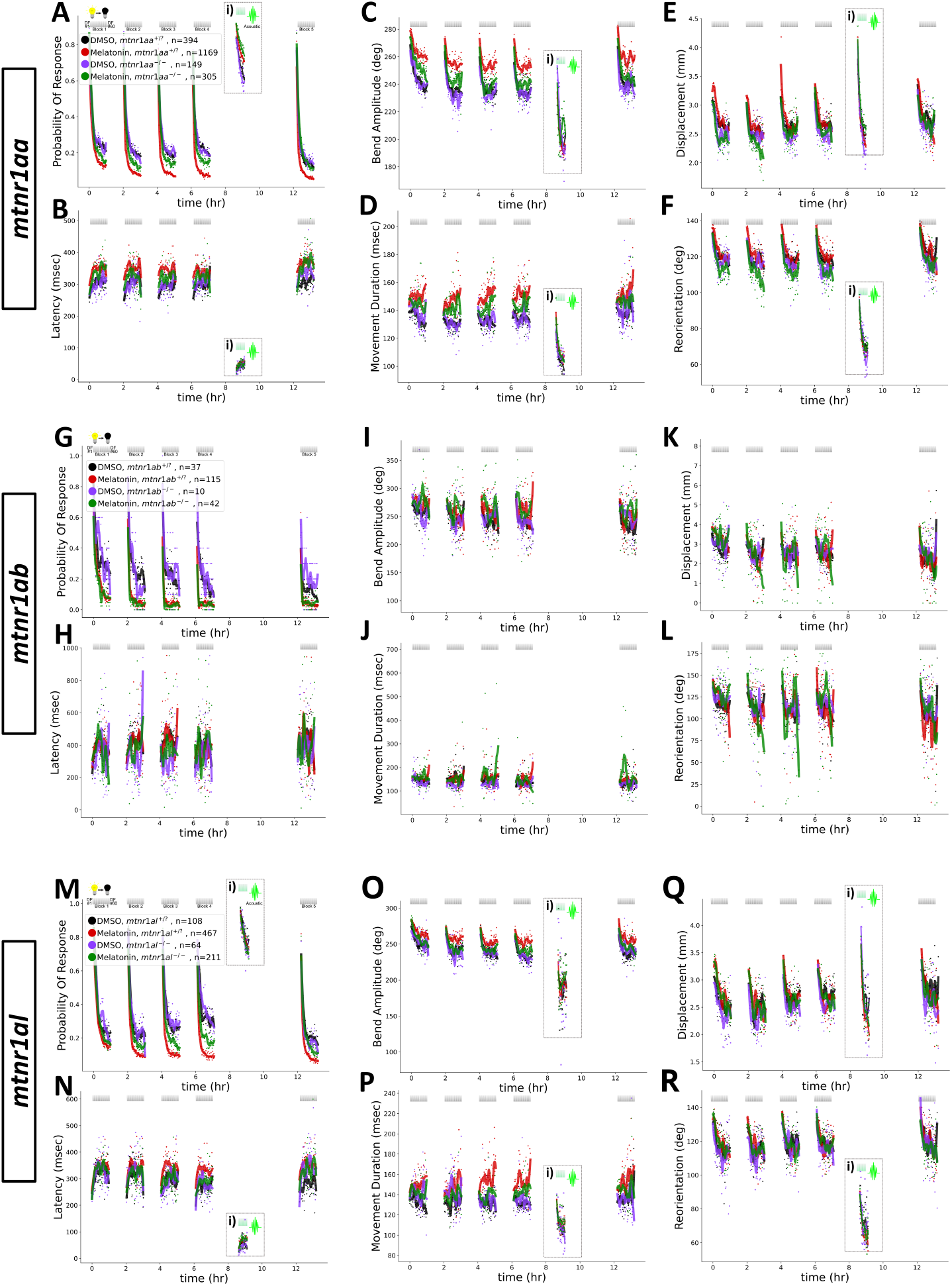
Zebrafish mutants for ***mtnr1aa*** and ***mtnr1al***, but not ***mtnr1ab***, are less sensitive to Melatonin’s effects on habituation learning Habituation behaviour comparing mutants for *mtnr1aa* (A-F; 9 different plates: DMSO *mtnr1aa^+/?^* n = 394, Melatonin *mtnr1aa^+/?^* n = 1169, DMSO *mtnr1aa^-/-^* n = 149, Melatonin *mtnr1aa^-/-^* n = 305), *mtnr1ab* (G-L; 2 different plates: DMSO *mtnr1ab^+/?^* n = 37, Melatonin *mtnr1ab^+/?^* n = 115, DMSO *mtnr1ab^-/-^* n = 10, Melatonin *mtnr1ab^-/-^* n = 42), and *mtnr1al* (M-R; 4 different plates: DMSO *mtnr1al^+/?^* n = 108, Melatonin *mtnr1al^+/?^* n = 467, DMSO *mtnr1al^-/-^* n = 64, Melatonin *mtnr1al^-/-^* n = 211) to their WT and heterozygous sibling controls. Larvae are treated with either 1 µM Melatonin, or vehicle (0.1% DMSO). DF stimuli are delivered at 1-minute intervals, in 4 blocks of 60 stimuli, separated by 1 hour of rest (from 0:00-7:00). 1.5 hours later a block of 30 vibration stimuli are delivered at 1-minute intervals (i). Each dot is the mean response across larvae to each DF, while ‘X’s mark mean responses to vibration stimuli. Lines are smoothed in time with a Savitzky–Golay filter (window = 15 stimuli, order = 2). Data is shown for the following behavioural components: A), G), M) the probability of response to the stimulus: B), H), N) the response latency, C), I), O) the maximum tail-bend amplitude achieved during the movement, D), J), P) the duration of the movement, E), K), Q) the displacement of the movement, and F), L), R) the reorientation achieved by the larvae.

Therefore, we next turned our attention to the other MT_1_-type receptors, Mtnr1ab and Mtnr1al. Mtnr1ab is the most closely related paralog to Mtnr1aa, but we found no evidence that *mtnr1ab* was necessary for the effect of Melatonin on habituation (**Figure 3G-L, *Figure S3G-L***). Melatonin-treated *mtnr1ab* mutants were indistinguishable from Melatonin-treated sibling controls for all behavioural components, indicating that Mtnr1ab is not required for the effects of Melatonin on habituation learning.

This left Mtnr1al as our next candidate. Indeed, we found that *mtnr1al* mutants showed a nearly identical phenotype to *mtnr1aa* mutants (**Figure 3M-R, *Figure S3M-R***). Mutants show a clear reduction in the effects of Melatonin on habituation learning for the probability of response (**Figure 3M**), and a similar trend for the other behavioural components that did not show a statistically significant difference, but where the response magnitude of Melatonin-treated mutants is intermediate between untreated- and Melatonin-treated-controls (**Figure 3N-R**). Therefore, we conclude that Mtnr1al is also partially required for the effects of Melatonin on habituation learning, having similar behavioural effects to Mtnr1aa.

#### Mtnr1ba, Mtnr1bb, and Mtnr1c are dispensable for the effects of Melatonin on habituation learning

Having identified two MT_1_-type receptors that contribute to mediating the effects of Melatonin on habituation learning, we next wanted to determine if the MT_2_-type receptors (Mtnr1ba and Mtnr1bb), or the third type of receptor (Mtnr1c), might also modulate habituation learning. As with *mtnr1ab*, we found no evidence that mutants for Mtnr1ba, Mtnr1bb, or Mtnr1c show deficits in the effects of Melatonin on habituation learning (**Figure 4, *Figure S4***). In each case, Melatonin-treated mutants were indistinguishable from Melatonin-treated controls, indicating that these receptors do not influence habituation learning.

**Figure 4.**
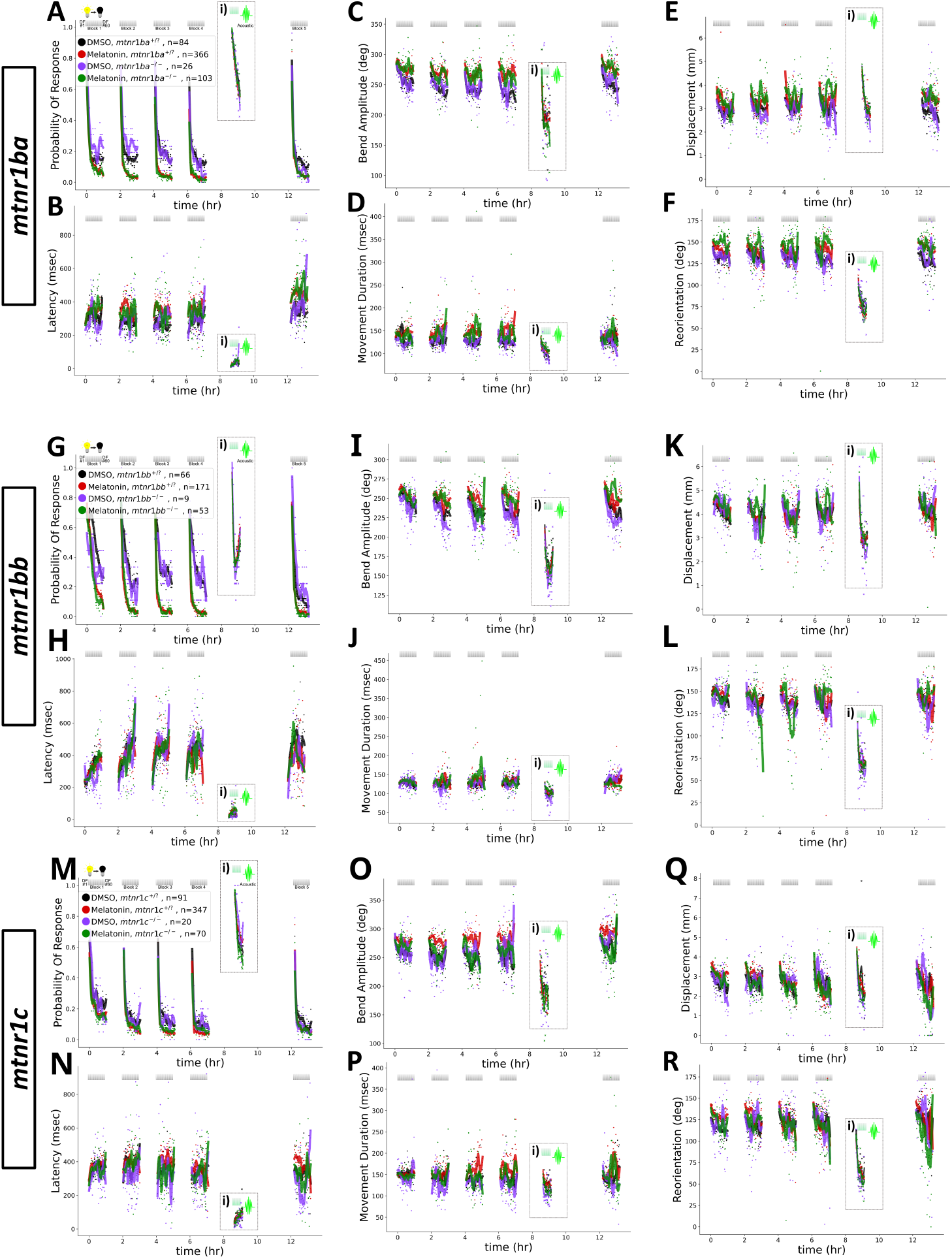
Zebrafish mutants for ***mtnr1ba*, *mtnr1bb***, and ***mtnr1c*** show normal responsiveness to Melatonin’s effects on habituation learning Habituation behaviour comparing mutants for *mtnr1ba* (A-F; 4 different plates: DMSO *mtnr1ba^+/?^* n = 84, Melatonin *mtnr1ba^+/?^* n = 366, DMSO *mtnr1ba^-/-^* n = 26, Melatonin *mtnr1ba^-/-^* n = 103), *mtnr1bb* (G-L; 1 plate: DMSO *mtnr1bb^+/?^* n = 66, Melatonin *mtnr1bb^+/?^* n = 171, DMSO *mtnr1bb^-/-^* n = 9, Melatonin *mtnr1bb^-/-^* n = 53), and *mtnr1c* (M-R; 2 different plates: DMSO *mtnr1c^+/?^* n = 91, Melatonin *mtnr1c^+/?^* n = 347, DMSO *mtnr1c^-/-^* n = 20, Melatonin *mtnr1c^-/-^* n = 70) to their WT and heterozygous sibling controls. Larvae are treated with either 1 µM Melatonin, or vehicle (0.1% DMSO). DF stimuli are delivered at 1-minute intervals, in 4 blocks of 60 stimuli, separated by 1 hour of rest (from 0:00-7:00). 1.5 hours later a block of 30 vibration stimuli are delivered at 1-minute intervals (i). Each dot is the mean response across larvae to each DF, while ‘X’s mark mean responses to vibration stimuli. Lines are smoothed in time with a Savitzky–Golay filter (window = 15 stimuli, order = 2). Data is shown for the following behavioural components: A), G), M) the probability of response to the stimulus: B), H), N) the response latency, C), I), O) the maximum tail-bend amplitude achieved during the movement, D), J), P) the duration of the movement, E), K), Q) the displacement of the movement, and F), L), R) the reorientation achieved by the larvae.

#### Mtnr1aa and Mtnr1al act together to mediate Melatonin’s effects on habituation learning

Our experiments with the individual mutants found that both Mtnr1aa and Mtnr1al contribute to mediating the effects of Melatonin on habituation learning, but that each contributes only partially, approximately half of the effect. Therefore, we wanted to determine if these receptors together can account for the full effects of Melatonin. To test this, we generated double mutants for *mtnr1aa* and *mtnr1al*, and tested their sensitivity to Melatonin. We found that the double mutants show a much more robust reduction in the effects of Melatonin on habituation learning for the probability of response (**Figure 5A, *Figure S5A***), where double mutants treated with Melatonin are significantly more responsive than Melatonin-treated sibling controls during all the habituation phases, and are indistinguishable from vehicle-treated sibling controls. This was also broadly true for the kinematic parameters modulated by Melatonin: latency, bend amplitude and movement duration. A significant statistical relationship was observed in most epochs we analyzed, and importantly always in the “Trained Response” epoch representing the period of training blocks 2-4 (***Figure S5Biv-Div***), where the Melatonin-dependent effect is the most robust (**Figure 1**). These results are consistent with a model in which the effects of Melatonin on habituation learning are mediated by both Mtnr1aa and Mtnr1al, leading to a complete loss of sensitivity to the effects of Melatonin on habituation learning in double mutants. Our data further indicate that this is true not only for the components of habituation that are potentiated by Melatonin (probability and latency), but also those that are inhibited (bend amplitude and movement duration), reflecting a cooperative relationship that is applied consistently across these different aspects of learning.

**Figure 5.**
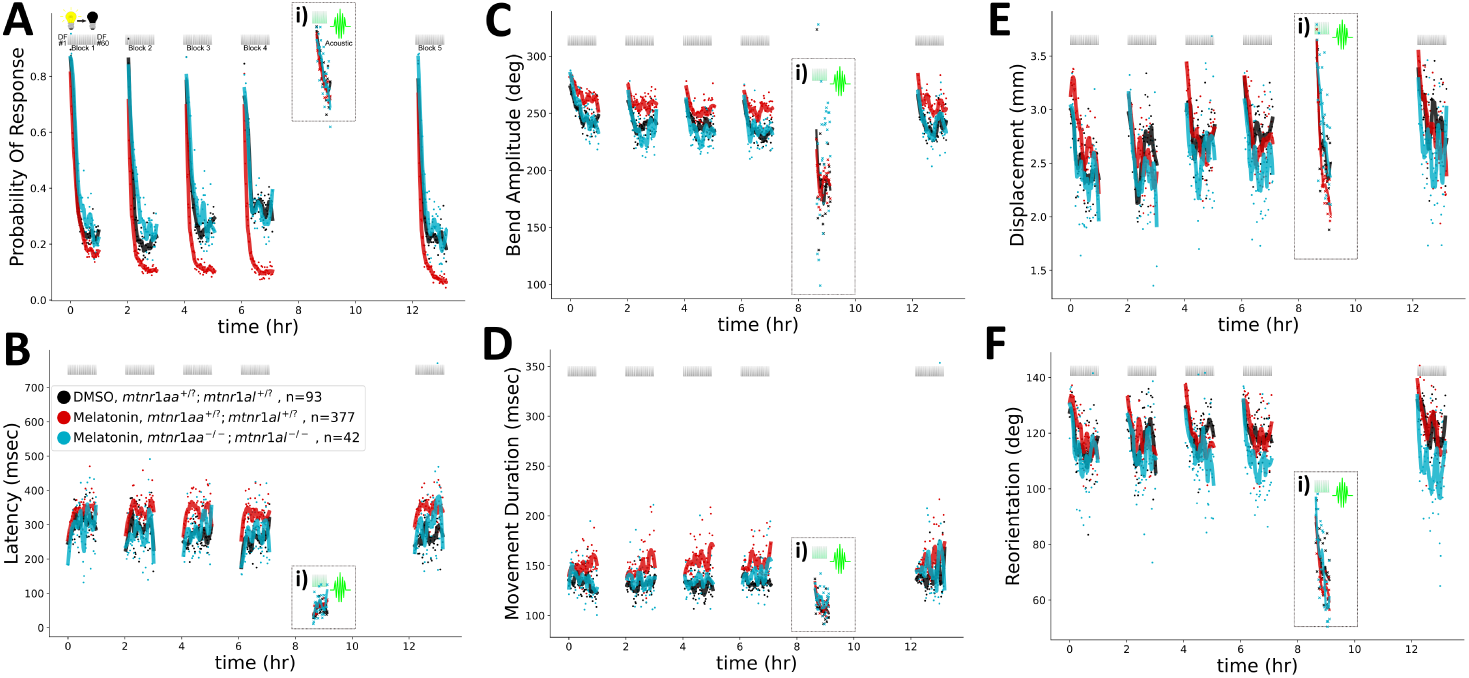
Zebrafish larvae lacking both ***mtnr1aa*** and ***mtnr1al*** are insensitive to Melatonin’s effects on habituation learning Habituation behaviour comparing double mutants for *mtnr1aa* and *mtnr1al* to their WT and heterozygous sibling controls (A-F; 3 different plates: sibling controls *mtnr1aa^+/?^; mtnr1al^+/?^* with DMSO n = 93 and Melatonin n = 377; double mutants *mtnr1aa^-/-^; mtnr1al^-/-^* with Melatonin n = 42). Larvae are treated with either 1 µM Melatonin, or vehicle (0.1% DMSO). Due to the rarity of the double mutants, we did not have enough double mutant larvae to test the vehicle treated group for analyses. DF stimuli are delivered at 1-minute intervals, in 4 blocks of 60 stimuli, separated by 1 hour of rest (from 0:00-7:00). 1.5 hours later a block of 30 vibration stimuli are delivered at 1-minute intervals (i). Each dot is the mean response across larvae to each DF, while ‘X’s mark mean responses to vibration stimuli. Lines are smoothed in time with a Savitzky–Golay filter (window = 15 stimuli, order = 2). Data is shown for the following behavioural components: A) the probability of response to the stimulus: B) the response latency, C) the maximum tail-bend amplitude achieved during the movement, D) the duration of the movement, E) the displacement of the movement, and F) the reorientation achieved by the larvae.

### Whole-brain activity mapping reveals distinct but partially overlapping activity patterns in ***mtnr1aa*** and ***mtnr1al*** mutants

Our behavioural experiments indicate that Mtnr1aa and Mtnr1al act together to mediate the effects of Melatonin on habituation learning, but it is unclear how these receptors functionally interact. We can envision two models for how Mtnr1aa and Mtnr1al could act in this context. In the first model, Mtnr1aa and Mtnr1al could be expressed in the same neurons, and therefore could act in a cooperative manner within the same cells, for example, each contributing half of the intracellular signalling effect. In the second model, Mtnr1aa and Mtnr1al could be expressed in different neurons, and therefore could act across different parts of the circuit to mediate the effects of Melatonin on habituation learning. To distinguish between these models, we first examined the expression pattern of Mtnr1aa and Mtnr1al in the brain using fluorescent *in situ* hybridization (HCR) for each of these receptors. This confirmed that Mtnr1aa is broadly expressed across the brain (Hill et al., 2026, **Figure 6A**), including in many brain regions that we have previously implicated in habituation learning using Ca^2+^ imaging (Lamiré et al., 2023), such as the tectum, tegmentum and cerebellum, as well as a cluster of neurons in the medial hindbrain, which appears to be a hotspot for Melatonin receptor-expression (Hill et al., 2026). We also observed clear *mtnr1aa* expression in the torus longitudinalis (TL) and preoptic area (POA) (**Figure 6A**), consistent with the *mtnr1aa* expression pattern reported by Hill et al. (2026). In contrast to *mtnr1aa*, they did not detect a clear signal for *mtnr1al*, but did see faint signal in the tectum. We observed some very sparse staining in the tectum for *mtnr1al*, but it was unclear to us if this staining might represent background signals. Therefore, while our expression data place Mtnr1aa at several candidate sites, they could not shed much light on where Mtnr1al might act.

**Figure 6.**
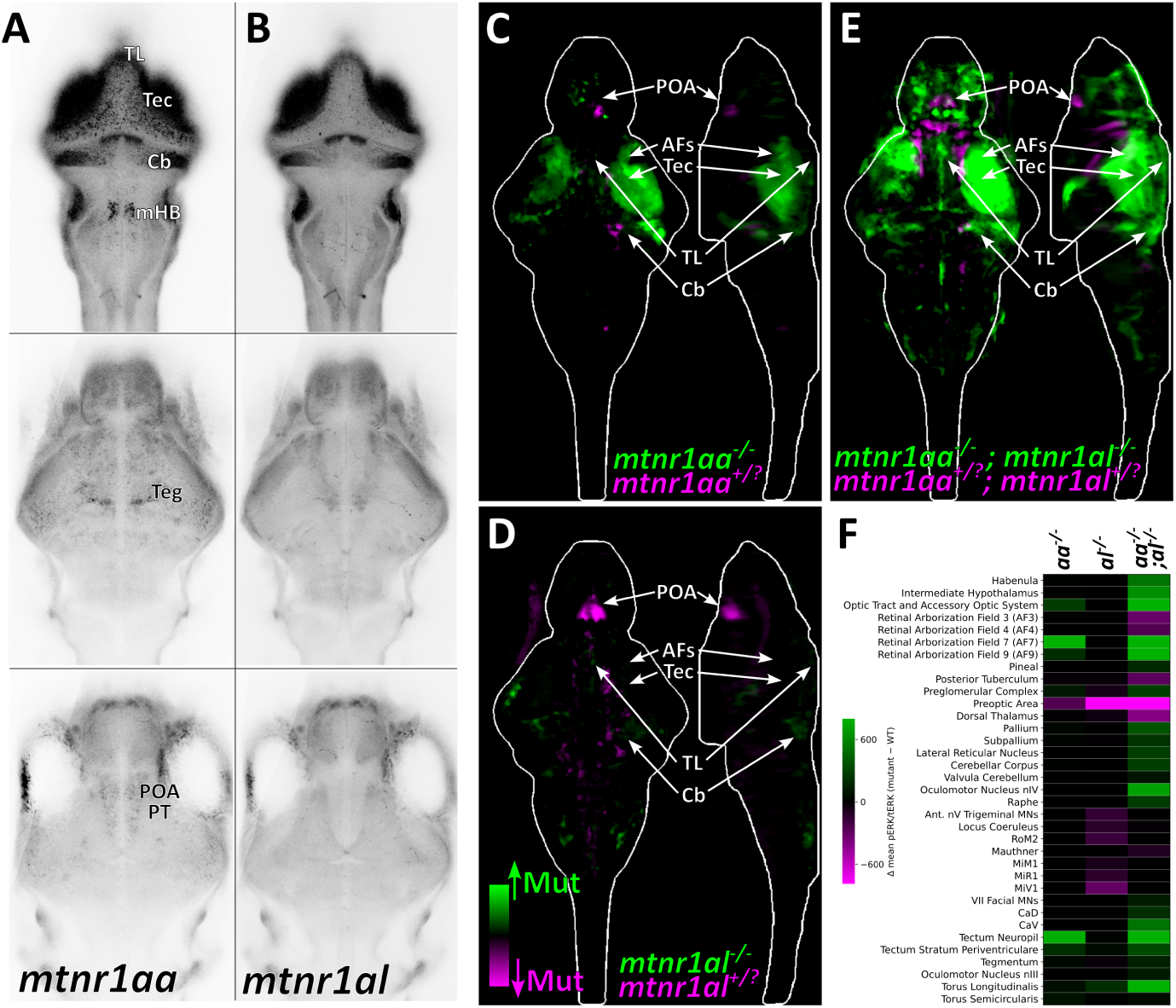
Circuit-level effects of ***mtnr1aa*** and ***mtnr1al*** during habituation learning A) Fluorescent *in situ* hybridization for *mtnr1aa* in the brain of 5dpf larval zebrafish. 3 planes of the brain are shown moving from dorsal (top) to ventral (bottom). Data are the average of 4 animals co-registered to the same coordinate space. Tec = tectum, Cb = Cerebellum, Teg = Tegmentum, TL = Torus Longitudinalis, POA = Preoptic Area, PT = Posterior Tuberculum, mHB = Medial Hindbrain. B) Fluorescent *in situ* hybridization for *mtnr1al*. Data are the average of 4 animals co-registered to the same coordinate space. C-E) pERK-based MAP-mapping of differential activity in mutants relative to sibling controls, both treated with Melatonin (1 µM) and stimulated with 60 DFs, in order to highlight mutation-dependent differences in neural activity during habituation, for: C) *mtnr1aa* mutants, D) *mtnr1al* mutants, and E) *mtnr1aa;mtnr1al* double mutants. Significant increases in the ratio of pERK/tERK staining in mutants relative to siblings are shown in green, while decreases are shown in magenta. All larvae were siblings from *mtnr1aa^+/-^;mtnr1al^+/-^* double-heterozygous incrosses, genotyped post-hoc, with each mutant class compared to the shared sibling control group (*mtnr1aa^+/?^;mtnr1al^+/?^*, n = 38): *mtnr1aa^-/-^;mtnr1al^+/?^* n = 19 (C), *mtnr1aa^+/?^;mtnr1al^-/-^* n = 20 (D), and *mtnr1aa^-/-^;mtnr1al^-/-^* n = 27 (E). F) Heatmap depicting the mean differential pERK/tERK in the different mutants from (C-E) in different brain areas of the Z-Brain atlas (MECE regions, see Vohra et al., 2025).

Since the expression data was inconclusive, we turned to a whole-brain functional mapping approach to try and identify the functional sites of action for these receptors. We used the pERK-based MAP-mapping method (Randlett et al., 2015) which maps differential activity comparing across different conditions (e.g. Mutant vs. WT). In order to identify where in the brain mutants show differential activity during habituation learning, we compared Melatonin-treated mutants vs. Melatonin-treated sibling controls after being exposed to a block of 60 dark flashes. This analysis revealed that the most prominent changes in the *mtnr1aa* mutants are in the tectum and the vicinity of the retinal arborization fields (AFs) which showed stronger activity in the mutants relative to sibling controls (**Figure 6C*, F***). This is consistent with a prominent role for the tectum, and perhaps other primary visual areas like the pretectum, in mediating habituation learning (Lamiré et al., 2023).

In contrast, *mtnr1al* mutants showed very little modulation of the tectum, indicating that this is likely not the primary site of action for this receptor. However, they showed very strong suppression of the preoptic area (POA) (**Figure 6D*, F***). The POA was also suppressed in the *mtnr1aa* mutants, but to a lesser extent. The torus longitudinalis (TL) was strongly activated in the *mtnr1al* mutants and, like the POA, showed weaker but consistent modulation in the *mtnr1aa* mutants. The *mtnr1al* mutants also showed sparse alterations in activity in distributed hindbrain regions, including the cerebellum, which was also increased in the *mtnr1aa* mutants.

To identify how these receptors function cooperatively, we analyzed double mutants for both *mtnr1aa* and *mtnr1al* (**Figure 6E*, F***). These double mutants showed the strongest differential activity pattern, consistent with the stronger behavioural effects observed in these mutants (**Figure 5**). Indeed, the double-mutant map combined the more localized effects seen in the individual mutants — activation of the tectum, retinal AFs, TL, and cerebellum, and suppression of the POA — together with additional changes in the pallium, subpallium, and habenula. The TL and cerebellar activation were both substantially further amplified in the double mutant relative to either single mutant, consistent with both receptors contributing additively to these shared nodes. The POA, though suppressed in both single mutants, was not more strongly suppressed in the double than in *mtnr1al* mutants alone, indicating that Mtnr1al may be the dominant receptor at this site. Collectively, these data support a model in which Mtnr1aa and Mtnr1al have unique effects across the brain, with Mtnr1aa having the more widespread effects, consistent with its broader expression pattern. However, the functional effects overlap and reinforce in some areas, providing a candidate anatomical basis for their non-redundant, cooperative control of habituation learning.

## Discussion

### Melatonin bidirectionally modulates distinct habituation mechanisms

A central conclusion from studies of habituation in larval zebrafish is that this behaviour is inherently modular: response probability, latency, amplitude, and duration each habituate through molecularly distinct and largely independent mechanisms operating in parallel across distributed brain circuits (Randlett et al., 2019; Nelson et al., 2023; Lamiré et al., 2023). This is consistent with the principle, established across organisms from *C. elegans* to mammals, that habituation is not a unitary plasticity process but emerges from multiple parallel plasticity events at distinct circuit nodes (Ramaswami, 2014; Cooke and Ramaswami, 2020; McDiarmid et al., 2019). Our data show that Melatonin engages this modular organization selectively and bidirectionally: it accelerates habituation learning for response probability and latency while slowing it for amplitude and duration, leaving displacement and reorientation largely unaffected. This is critical for our interpretation of the action of Melatonin, which is best known as a promoter of sleep, and therefore it is important to establish whether behavioural alterations induced by Melatonin are simply byproducts of a change in arousal or state. The bidirectional pattern we observe would seem to be incompatible with a global arousal-suppressing action — if Melatonin were simply inducing sedation, we would expect all response parameters would shift in the sluggish direction. Additionally, probability of responding to an acoustic stimulus is indistinguishable between Melatonin-treated and control animals at the concentrations used (**Figure 1Ai**, Lamiré et al., 2023), indicating that Melatonin’s effects are sensorimotor-circuit-specific.

Recently, Hill et al. (2026) propose that Melatonin acts to suppress visual responses, and that this mechanism of stimulus insensitivity may explain the sleep-promoting effects of Melatonin. They found that Melatonin-treated larvae were less sensitive to a weak visual dimming stimulus when that stimulus followed a strong dimming stimulus that elicited a response. Because the dimming stimulus was presented 20 s after a strong dark flash and behaviour was scored as a binary response, this reduction could reflect Melatonin-dependent modulation of short-term adaptation to, or integration of, successive visual stimuli. Therefore, we propose that Melatonin regulates visual behaviour by modulating plasticity and adaptation to recent stimulus history, rather than simply suppressing responsiveness.

The bidirectional nature of Melatonin’s effects on plasticity is not without precedent. Exogenous Melatonin modulates multiple distinct forms of synaptic plasticity in hippocampal slices in a parameter-dependent manner, attenuating low-frequency evoked population spikes while modifying paired-pulse facilitation and inhibition in opposite directions depending on inter-stimulus interval (El-Sherif et al., 2003). Melatonin’s effects on LTP itself are also complex: MT_2_ receptor signalling is required for normal hippocampal LTP (Larson et al., 2006), implying Melatonin promotes this form of potentiation, yet loss of *both* MT_1_ and MT_2_ receptors paradoxically enhances cognitive and motor performance (O’Neal-Moffitt et al., 2014) — indicating that Melatonin signalling simultaneously constrains plasticity in other contexts. At the behavioural level, individual receptor knockouts impair specific forms of memory (Pistono et al., 2021; Clough et al., 2014), while in zebrafish exogenous Melatonin suppresses nighttime operant memory formation (Rawashdeh et al., 2007) but potentiates visual habituation in our hands. Thus the sign of Melatonin’s effect on plasticity is determined by paradigm, circuit, and receptor context rather than reflecting a fixed pro- or anti-plasticity role. At the molecular level, Gi-coupled Melatonin receptor signaling converges on cAMP/CREB and downstream kinase pathways whose consequences for synaptic plasticity are highly state- and circuit-dependent (Valdés-Tovar et al., 2018; Jilg et al., 2019; Malenka and Bear, 2004). The specific context of habituation training — where distinct molecular programs independently regulate distinct response components — thus provides a revealing readout through which the bidirectional circuit-specificity of Melatonin can be dissected.

### The role of endogenously produced Melatonin in habituation learning

We observed a surprisingly subtle effect when we studied the habituation behaviour of Melatonin deficient *aanat1;aanat2* double mutants, where they exhibit a weak deficit only for the response probability, but not the other behavioural parameters modulated by exogenously applied Melatonin. We believe that this may reflect the fundamental limitations of the visual assay: dark-flash stimulation requires light-adaptation, which is itself a potent suppressor of pineal and retinal Melatonin synthesis. The more selective deficit of endogenous deficiency compared to the bidirectional effects of pharmacological Melatonin may therefore reflect a physiological ceiling on achievable Melatonin concentrations during the assay, rather than a true mechanistic difference in the circuits that endogenous Melatonin normally modulates.

A notable feature of these experiments is that the habituation deficit requires loss of both Aanat1 and Aanat2, with neither single mutant alone producing a phenotype. This functional redundancy is interesting in light of the distinct anatomical sources of these two enzymes: Aanat2 drives Melatonin synthesis in both the pineal gland and the retina, whereas Aanat1 is expressed almost exclusively in the retina. Previous behavioural studies of *aanat* mutant larvae have focused primarily on the pineal gland as the relevant source of systemic hormone: Gandhi et al. (2015) found that *aanat2* single mutants show disrupted sleep — consistent with a dominant pineal contribution to circulating nocturnal Melatonin — while *aanat1* single mutants did not have a phenotype in this context. Similarly, Rawashdeh et al. (2007) attributed the night-time suppression of operant memory to pineal-derived Melatonin, based on the effects of pinealectomy. Hill et al. (2026) demonstrated that endogenous Melatonin suppresses visual responsiveness at night through *aanat2*-derived Melatonin, explicitly noting that their *aanat2* mutants retain retinal Melatonin (from Aanat1), and separately confirming that *aanat1* mutants do not worsen the *aanat2* sleep phenotype — establishing pineal-derived Melatonin as the primary relevant source in that context. Our habituation data suggest a different relationship between these two Melatonin sources: the requirement for loss of *both* Aanat1 and Aanat2 demonstrates that either source alone is sufficient to sustain normal habituation. An interesting implication of this redundancy is that the pineal gland may not be the dominant source of Melatonin for habituation learning, and that retinal Melatonin (derived from Aanat1 or Aanat2) may be sufficient to sustain normal habituation in the absence of pineal-derived Melatonin in *aanat2* mutants. Consistent with this interpretation, we observed no circadian dependence of the habituation deficit — day-tested (**Figure 2A**) and night-tested (**Figure 2G**) double mutants showed similarly impaired habituation.

Together, these observations raise the possibility that retinal Melatonin — produced locally in a light-regulated, cell-autonomous manner — is the dominant endogenous source modulating habituation in this context, with the two retinal AANAT paralogs providing redundant synthetic capacity. Retinal Melatonin is generally considered a local neuromodulator, acting to regulate photoreceptor function and retinal signal processing (Besharse and McMahon, 2016; Bhoi et al., 2023), and therefore it could exert its influence on habituation via the modulation of retinal ganglion cell activity. In the adult fish retina, Melatonin acts through Melatonin receptors and the dopaminergic system to modulate retinal signalling — reducing the responsiveness of cone-driven horizontal cells and shifting the balance between cone and rod pathways (Ribelayga et al., 2004; Huang et al., 2012). The larval zebrafish retina is cone-dominated at the stages we study, with rod-driven vision maturing only later (Saszik et al., 1999), so the rod-related components of this switch are unlikely to apply; nonetheless, Melatonin’s modulation of cone-driven circuitry and retinal dopamine provides a plausible route by which endogenous Melatonin could bias the visual drive feeding the dark-flash habituation circuit. However, Melatonin is a small (∼232 Da), highly lipophilic molecule that crosses membranes by passive diffusion. Melatonin produced in the retina could conceivably diffuse inward through the retina, and along the optic nerve towards central targets. Notably, some of the most prominent sites of differential pERK activity in mutants are proximal to the termination zones of retinal ganglion cell axons, including the tectum and the preoptic area. Whether Mtnr1aa/l at these locations are activated by retinal Melatonin arriving by diffusion, by systemic Melatonin, or by Melatonin modulating the retinal output signal itself remains to be determined, but the anatomical coincidence between the site of Melatonin receptor action and the termination of retinal projections makes the retina a compelling candidate source.

### Functional cooperation of two Melatonin receptors

Of the six zebrafish Melatonin receptors, only the two MT_1_-type paralogs Mtnr1aa and Mtnr1al contribute to Melatonin’s effects on habituation. In mammals, MT_2_ is required for normal hippocampal long-term potentiation (Larson et al., 2006) and long-term recognition memory (Pistono et al., 2021), and deletion of either MT_1_ or MT_2_ alone is sufficient to abrogate methamphetamine-induced conditioned place preference (Clough et al., 2014), demonstrating functional roles for both subtypes in plasticity. Therefore, it is clear that MT_2_-type receptors are able to influence plasticity in some contexts. While this may highlight a difference between zebrafish and mammalian MT_2_-type receptors, a more likely explanation for their dispensability in our assay is that they may simply not be expressed in the relevant circuits implementing habituation. Differential expression may also explain the dispensability of the MT_1_-type receptor Mtnr1ab, as well as Mtnr1c.

The non-redundancy of Mtnr1aa and Mtnr1al adds a further layer of specialization: each single mutant loses approximately half of the Melatonin-dependent habituation enhancement across all response components, and double mutants lose it entirely. This cooperative relationship between two receptors that reinforce a common outcome may represent a more general and more modular strategy for hormonal control of behaviour, where the signal is distributed across parallel channels, rather than duplicated within a redundant pair.

### Circuit-level action of Melatonin receptors during habituation

Our whole-brain pERK activity mapping revealed that the two receptors have very distinct functional effects on brain activity, that nonetheless converge on shared nodes: the torus longitudinalis (TL), cerebellum, and preoptic area (POA). In Melatonin-treated *mtnr1aa* mutants, the optic tectum and retinal arborization fields show the most prominent increases in activity after dark-flash training relative to sibling controls (Randlett et al., 2019; Lamiré et al., 2023). Mtnr1al mutants, by comparison, show minimal tectal modulation, but the POA and TL are perturbed in both single mutants — the POA suppressed and the TL activated — strongly in *mtnr1al* mutants and more weakly in *mtnr1aa* mutants. In double mutants these changes are combined, and the TL showed clear amplification relative to either single mutant, consistent with both receptors contributing to this shared node.

The POA is a conserved vertebrate hub for sleep and arousal, where galanin-expressing neurons integrate sleep pressure (Reichert et al., 2019) and MT_1_/MT_2_ receptors have been proposed to gate sleep onset (Ng et al., 2017; Klosen et al., 2019); in our context, Mtnr1aa and Mtnr1al appear to sustain POA activity during training. Classical dual-process accounts of habituation hold that behavioural output reflects not only a stimulus-specific decremental process but also a separable state/arousal system that sets overall excitability (Groves and Thompson, 1970); the POA is a natural candidate substrate for such a state system, providing a route by which a hormonal signal like Melatonin could shape habituation output through the arousal axis rather than through the stimulus-specific plasticity itself. Alternatively, POA-dependent modulation could feed more directly into the visual habituation system through specific neuropeptidergic outputs or through connectivity with visual sensorimotor circuits that set response gain during training.

The TL is a midbrain structure that provides glutamatergic input to apical dendrites of tectal pyramidal neurons (Folgueira et al., 2020; DeMarco et al., 2021), is required for binocularity and spatial summation of visual responses in the tectum (Tesmer et al., 2022), and participates in a dimming-selective feedback pathway from the tectum (Robles et al., 2021) engaged by our dark-flash stimulus. Consistent with a role in this stimulus regime, TL projection neurons respond most strongly to dimming when it follows low ambient light (Robles et al., 2021), and sustained TL activity is evoked by dark stimuli (Northmore, 2017); the TL has further been proposed to prime tectal pyramidal-cell responses through connections that could be shaped by experience (Northmore, 2017). It is therefore well-placed for a modulator like Melatonin to bias the responses to dark flashes and, in turn, the trajectory of their habituation.

The cerebellum showed the same pattern as the TL — activated in both single mutants and further amplified in the double — and therefore represents another potential shared convergence node for the two receptors. The cerebellum has been implicated in motor-related learning in larval zebrafish (Markov et al., 2021; Lin et al., 2020), including behaviours driven by visual error signals relayed through the inferior olive (Ahrens et al., 2012). Our previous Ca^2+^ imaging is also consistent with an important role for the cerebellum in habituation, where we identified interesting neuronal classes within this structure, including non-adapting and potentiating neuronal populations: populations that maintain or increase their activity as habituation proceeds, respectively (Lamiré et al., 2023). Melatonin, acting via Mtnr1aa and Mtnr1al expressed in or projecting to the cerebellum, could therefore modulate DF habituation via the cerebellum, in parallel with effects on visual gain at the TL and arousal state at the POA.

A notable feature of this relationship is that both Mtnr1aa and Mtnr1al appear to contribute equivalently to all behavioural components modulated by Melatonin — probability, latency, amplitude, and duration — rather than each selectively modulating a distinct subset of habituation parameters. This is perhaps surprising given the model in which each response component is governed by largely independent plasticity mechanisms operating in parallel (Randlett et al., 2019; Nelson et al., 2023). More speculatively, the different nodes may map onto the different directions of Melatonin’s bidirectional effect. For example, Melatonin suppresses TL activity, and because the TL sets the visual gain that scales the vigour of the tectal-driven response, this suppression could maintain movement amplitude and duration against habituation — the *inhibited* arm. Melatonin also sustains POA activity, and as a conserved arousal hub the POA is well placed to drive the *potentiated* arm — the enhanced habituation of response probability and latency. In this scheme the direction of modulation is set by which node is engaged, while both Mtnr1aa and Mtnr1al perturb both nodes, with Mtnr1al dominating at the POA and both receptors contributing additively at the TL; this would explain why loss of either receptor blunts both directions of the effect, rather than selectively removing one. This mapping is speculative, but it makes a concrete, testable prediction: that selectively manipulating POA versus TL activity should dissociate the probability and amplitude arms of visual habituation.

An important caveat for these analyses is that differential pERK identifies where neural activity is altered in the mutants, not necessarily where the receptors act directly. The strongest pERK changes could reflect secondary consequences of receptor action upstream: for example, elevated tectal pERK in *mtnr1aa* mutants could result from Mtnr1aa acting on tectal neurons, but equally from Mtnr1aa acting upstream in the retina, or on presynaptic inputs to the tectum. Similarly, whether the POA and TL changes in mutants reflect independent primary sites of receptor action, or whether one is a downstream consequence of the other, or of a site elsewhere, cannot be resolved from these activity maps alone.

## Conclusions

We find that two MT_1_-type paralogs, Mtnr1aa and Mtnr1al, cooperatively and non-redundantly mediate Melatonin’s bidirectional control of habituation learning, and propose that they do so by acting at partly distinct sites that converge on shared circuit nodes, including potentially the TL, cerebellum, and POA. This contrasts with the redundant logic these same receptors follow in sleep regulation (Hill et al., 2026), illustrating how an expanded teleost receptor repertoire can be deployed with different genetic logic to parcellate modulatory influence with finer control. Resolving the synaptic mechanisms through which these receptors tune the multiple plasticity streams that control visual habituation will require cell-type-specific imaging together with chemogenetic or optogenetic perturbation of POA, TL, cerebellar, and tectal neurons in animals expressing or lacking each receptor.

## Materials and Methods

### Animal husbandry and care

All experiments were carried out using larval zebrafish at 5 days post-fertilization (dpf), maintained in E3 embryo medium supplemented with 0.02% HEPES (pH 7.2) at a density of approximately 1 larva per mL. Larvae were housed in 10 cm petri dishes under a 14:10 h light/dark cycle (lights-on at 08:00, lights-off at 20:00) at a constant temperature of 28-29°C. Zebrafish were maintained and bred at two facilities: the Plateau de Recherche Expérimentale en Criblage In Vivo (PRECI, SFR Biosciences, Lyon) and the Animalerie Zebrafish Rockefeller (AZR, SFR Santé Lyon Est, Lyon). Adult zebrafish husbandry was carried out in compliance with the regulations of the PRECI and AZR facilities, under the oversight of the regional ethics committee (comité d’éthique en expérimentation animale de la Région Rhône-Alpes: CECCAPP, Agreement #C693870602). Larval behavioural experiments were conducted at 5 dpf and are therefore not subject to ethical review under French regulations. Mutants for the Melatonin-synthesizing arylalkylamine N-acetyltransferase genes (*aanat1*, *aanat2*) were a gift from Dr David Prober, and are as described in Gandhi et al. (2015).

### Generation of CRISPR mutant lines

CRISPR-Cas9 mutant lines targeting all six zebrafish Melatonin receptor genes (*mtnr1aa*, *mtnr1ab*, *mtnr1al*, *mtnr1ba*, *mtnr1bb*, *mtnr1c*) were generated using standard zebrafish CRISPR injection protocols in the TLF background (ZDB-GENO-990623-2). Guide RNAs (gRNAs) were designed to target early coding exons of each gene using the Millipore CRISPR Design Tools, and were synthesized by Sigma. Ribonucleoprotein complexes were assembled by combining gRNA with recombinant Cas9 protein (Sigma) and injected into the cell of one-cell-stage embryos. Founders (F0) were outcrossed to wild-type fish, and heterozygous F1 carriers were identified by PCR followed by Sanger sequencing of the target locus from fin-clip genomic DNA. For each gene, an allele bearing a frameshift-inducing insertion or deletion in the target area was selected and propagated. gRNA sequences and resulting indel mutations for each line are provided in ***Table 1***. Lines were maintained as heterozygous stocks. For behavioural experiments, clutches were generated from heterozygous incrosses, and larvae were genotyped after behavioural testing, such that all genotype classes were tested together within the same plates and the experimenter was blind to genotype during data acquisition; homozygous mutants were compared with pooled heterozygous and wild-type siblings, as we never observed any phenotypic differences between heterozygotes and wild-type siblings. The *mtnr1aa^-/-^*;*mtnr1al^-/-^* double-mutant line was generated by intercrossing the double-mutant heterozygous line *mtnr1aa^+/-^*;*mtnr1al^+/-^*.

### Genotyping

Larvae were individually lysed in 50 mM NaOH at 95°C for 10 min and neutralized with 1 M Tris-HCl (pH 8.0) at a 1:9 ratio (v/v) to extract genomic DNA. Target loci were amplified using FIREPol® Master Mix Ready to Load 1× (Solis Biodyne) with gene-specific primers at 0.1 µM. Alleles with resolvable size differences were distinguished by fragment analysis on high-density (2.5–3%) agarose gels. Primer sequences and expected amplicon sizes for each mutant line are provided in Supplementary Table 1.

### Behavioural testing and drug treatments

Melatonin (Sigma-Aldrich, M5250) was dissolved in DMSO to a 1 mM (1000x) stock solution and stored at −20°C. Working solutions were prepared fresh before each experiment by a 1:100 dilution into E3 yielding a 10 µM (10x) Melatonin stock solution in 1% DMSO. Larvae were distributed into the 300-well plates with one larva per well in 225 µL of E3 using a p1000 pipette tip with the end cut off to allow for gentle transfer. 25 µL of the 10 µM Melatonin stock was then added directly into the wells of the 300-well plate containing larvae, yielding a final concentration of 1 µM Melatonin in 0.1% DMSO. Vehicle controls consisted of E3 containing 0.1% DMSO, via the same dilution procedure. Experiments were generally started in the morning/early afternoon, between 11:00-13:00. For experiments using *aanat* double mutants, larvae were maintained in drug-free E3 throughout. To test Melatonin responses at circadian night, *aanat1^-/-^;aanat2^-/-^* larvae were raised under a reversed 14:10 light/dark cycle such that lights-off coincided with the start of the experimental period (11:00); behavioural testing was performed during this reversed dark phase, approximately at ZT12 of the reversed cycle. For these *aanat* mutant experiments, larvae were loaded into the plate at 4dpf and kept in the incubator overnight until testing at 5dpf, to minimize handling and light exposure on the testing day.

### Analyses of the Dark Flash Response

Larval behaviour was assessed in custom 300-well acrylic plates (8 mm diameter, 6 mm depth; ∼250 µL water volume) as previously described (Hsiao et al., 2025). The plates were positioned beneath a 31°C water bath, which served as a heated lid to prevent condensation and maintain the wells at 29°C. Recordings were captured using a Mikrotron CXP-4 camera (500 Hz) connected to a Silicon Software frame grabber (Marathon ACX-QP, Basler) and illuminated with IR LEDs (TSHF5410, digikey.com). Visual stimulation was provided by a rectangular array of 155 WS2813 RGB LEDs, with dark flashes (DFs) consisting of a 1 second light-off period followed by a linear return to baseline over 19 seconds. Acoustic/vibration taps were delivered via a solenoid (ROB-10391, Sparkfun). The experimental design alternated 1 hour of stimulation with 1 hour of rest, during which a stepper motor-driven actuator (Hanpose HPV4, 500 cm) moved the camera between two plates, enabling up to 600 larvae to be tested simultaneously.

Control of the LEDs, solenoid, and camera actuator was managed by a Raspberry Pi Pico running CircuitPython and custom Python software, which also handled video acquisition and online tracking of head and tail positions at 20-30 Hz. During DF or tap delivery, a 1 second high-speed burst recording was captured and later analyzed offline using background subtraction and morphological operations. Tracking and analysis relied on open-source libraries, including OpenCV, scikit-image, NumPy, SciPy, and Numba.

### Fluorescent ***in situ*** hybridization (HCR)

To map the expression of *mtnr1aa* and *mtnr1al* in the larval brain, we performed whole-mount fluorescent *in situ* hybridization (HCR v3.0) using probes designed by Molecular Instruments (Los Angeles, CA), using their standard protocol (Choi et al., 2018), and imaged from the dorsal surface on a Zeiss 880 confocal microscope using a 10× air objective with NA = 0.45 (CIQLE imaging platform, Lyon France).

### MAP-Mapping Analyses of Brain-Wide Activity Patterns

Whole-brain pERK activity mapping was performed using the MAP-mapping approach (Randlett et al., 2015). Larvae (5dpf) from crosses of heterozygous Melatonin receptor double mutants were treated with 1 µM Melatonin. Approximately one hour after treatment onset, larvae were transferred in groups to 10 cm petri dishes in the dark flash behaviour setup and allowed to acclimatize for 20 minutes. Larvae were then subjected to a block of 60 dark flashes (1 second light-off, 19 second linear ramp to lights-on, 1 minute inter-stimulus interval) prior to immediate fixation in 4% paraformaldehyde in PBS. Fixed larvae were processed for immunohistochemistry using anti-phospho-p44/42 MAPK (ERK1/2) antibody for pERK (Cell Signaling Technology, #4370S) and anti-p44/42 MAPK (ERK1/2) antibody for total ERK (tERK; Cell Signaling Technology, #4696S), followed by fluorophore-conjugated secondary antibodies: Alexa Fluor™ 647 for tERK (Thermo Fisher Scientific, #A21235) and Alexa Fluor™ 568 for pERK (Thermo Fisher Scientific, #11036). Brains were imaged on a Zeiss 880 confocal microscope using a 10× air objective with NA = 0.45 (CIQLE imaging platform, Lyon France). Image stacks were acquired at a voxel resolution of 0.08 × 0.08 × 2 µm, and two partially overlapping volumes were tiled to encompass the entire brain. The tERK channel was used as an anatomical reference for nonlinear registration of each specimen to a standardized larval zebrafish reference brain via the Computational Morphometry Toolkit (CMTK). Registered image volumes were subsequently down-sampled to 300 × 679 × 80 voxels (x, y, z) and spatially smoothed using a two-dimensional Gaussian filter. To account for inter-individual variability in staining intensity, voxel-wise pERK signals were normalized to corresponding tERK values prior to statistical comparison across experimental groups. Each mutant genotype was analyzed independently, comparing homozygous mutant larvae to pooled sibling controls (heterozygotes and wild-type) within the same Melatonin treatment condition. Larvae were derived from *mtnr1aa^+/-^*;*mtnr1al^+/-^* double-heterozygous incrosses and sorted post-hoc into genotype classes, such that all groups shared the same clutches, plates, and control cohort: *mtnr1aa^-/-^;mtnr1al^+/?^* (n = 19), *mtnr1aa^+/?^;mtnr1al^-/-^* (n = 20), *mtnr1aa^-/-^;mtnr1al^-/-^* (n = 27), and *mtnr1aa^+/?^;mtnr1al^+/?^* sibling controls (n = 38). Genotypes were confirmed post-hoc from tissue collected after brain imaging. Voxel-wise statistical comparisons and thresholding were performed as described (Randlett et al., 2015). These analyses were performed using custom Python scripts, available at github.com/owenrandlett/2026_MtnRs_Hab/code/MAPMapping.

## Data analysis and statistics

All behavioural data analyses were performed using custom Python scripts (NumPy, SciPy, Pandas, Matplotlib, Seaborn, scikit-posthocs), available at github.com/owenrandlett/2026_MtnRs_Hab/code. Per-fish group-average response traces across the habituation session were smoothed using a Savitzky-Golay filter (window = 15 stimuli, polynomial order = 2) for visualization. For statistical analysis, responses were averaged within each defined epoch per fish. Epoch means were plotted as individual data points (jittered strip plot) overlaid on violin plots; the violin shape represents the kernel density estimate of the per-fish epoch means, using a Gaussian kernel with Scott’s bandwidth rule and restricted to the range of the data (cut=0). Group means are indicated by white circles. Statistical comparisons between two groups were performed using the Mann-Whitney U test. For comparisons among three or more groups, the Kruskal-Wallis H-test was used, followed, where the Kruskal-Wallis test was significant, by Dunn’s post-hoc pairwise comparisons with Bonferroni correction for multiple comparisons, as implemented in the scikit-posthocs library (Terpilowski, 2019).

## Data and code availability

All code used to acquire and analyse the behavioural and imaging data in this study is available at github.com/owenrandlett/2026_MtnRs_Hab. Links to the underlying behavioural and imaging datasets are provided in the repository.

## Acknowledgments

We are grateful to the staff of the PRECI and AZR zebrafish facilities, including Laure Benard, Robert Renard, Annie Desenfant, Olivier Lohez, and Marion Delous for the expert care provided to the zebrafish. Imaging was performed in the CIQLE imaging platform (LYMIC, Lyon), and we thank the staff of the CIQLE, incuding Denis Ressnikoff and Bruno Chapuis for their support with confocal microscopy and image processing. We also gratefully acknowledge the communities that develop and maintain the numerous open-source software packages we rely on, most of which could not be cited here.

## Funding

This work was supported by funding from the ATIP-Avenir program of the CNRS and Inserm, a Fondation Fyssen research grant, and the IDEX-Impulsion initiative of the University of Lyon.

## Competing interests

The authors declare no competing interests.

---

**Figure S1.**
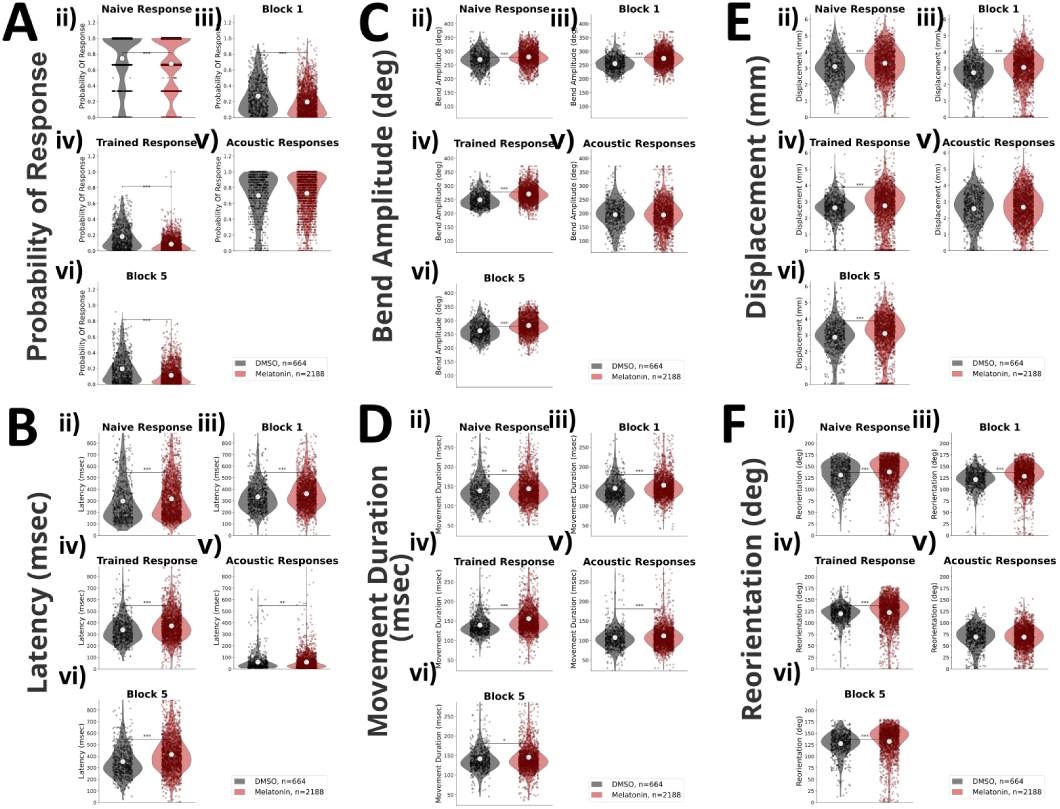
Quantification of epoch-level behavioural responses from Figure 1 Distributions of responsiveness for different epochs of the experiment ii)-vi): ii) Naive Response (first 3 dark-flash stimuli), iii) Block 1 (remaining 57 stimuli of the first training block), iv) Trained Response (training blocks 2–4), v) Acoustic Response (30 vibration stimuli), and vi) Re-test Block (Block 5, 5 hours after the final training block). Each dot is the per-fish average of the epoch. Statistical significance was calculated using the Mann-Whitney U test, * = p < 0.05, ** = p < 0.01, *** = p < 0.001. A) The probability of response to the stimulus, B) the response latency, C) the maximum tail-bend amplitude achieved during the movement, D) the duration of the movement, E) the displacement of the movement, and F) the reorientation achieved by the larvae.

**Figure S2.**
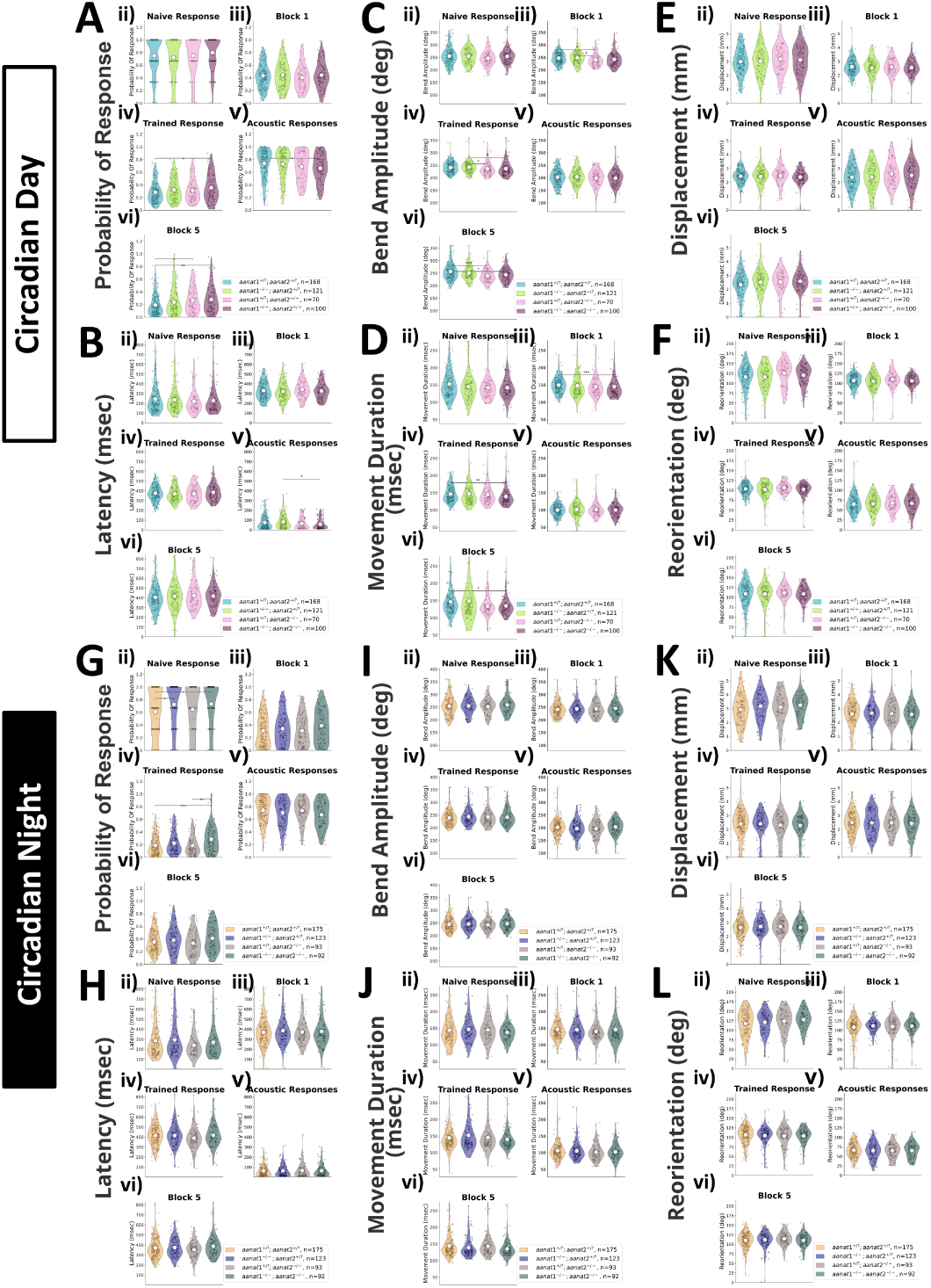
Quantification of epoch-level behavioural responses from Figure 2. Distributions of responsiveness for different epochs of the experiment ii)-vi): ii) Naive Response (first 3 dark-flash stimuli), iii) Block 1 (remaining 57 stimuli of the first training block), iv) Trained Response (training blocks 2–4), v) Acoustic Response (30 vibration stimuli), and vi) Re-test Block (Block 5, 5 hours after the final training block). Each dot is the per-fish average of the epoch. Statistical significance was calculated using the Kruskal-Wallis test followed by Dunn’s test for post-hoc pairwise comparisons with Bonferroni correction, * = p < 0.05, ** = p < 0.01, *** = p < 0.001. A-F) Day-tested cohort (ZT4), with A) the probability of response, B) the response latency, C) the maximum tail-bend amplitude, D) the duration, E) the displacement, and F) the reorientation. G-L) Night-tested cohort (shifted light cycle), with G) the probability of response, H) the response latency, I) the maximum tail-bend amplitude, J) the duration, K) the displacement, and L) the reorientation.

**Figure S3.**
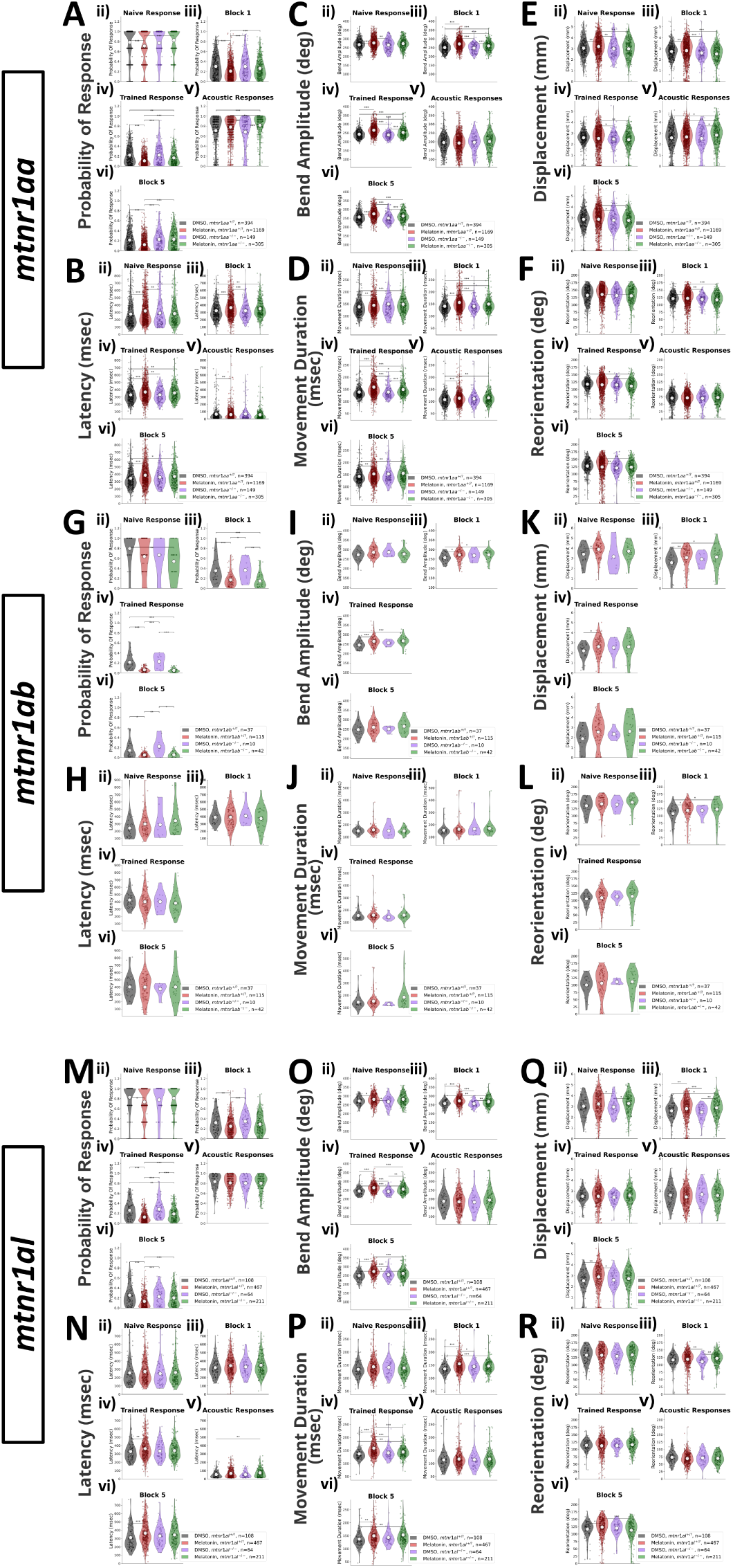
Quantification of epoch-level behavioural responses from Figure 3. Distributions of responsiveness for different epochs of the experiment ii)-v): ii) Naive Response (first 3 dark-flash stimuli), iii) Block 1 (remaining 57 stimuli of the first training block), iv) Trained Response (training blocks 2–4), and v) Acoustic Response (30 vibration stimuli). Each dot is the per-fish average of the epoch. Statistical significance was calculated using the Kruskal-Wallis test followed by Dunn’s test for post-hoc pairwise comparisons with Bonferroni correction, * = p < 0.05, ** = p < 0.01, *** = p < 0.001. A-F) *mtnr1aa* cohort, with A) probability of response, B) response latency, C) maximum tail-bend amplitude, D) duration, E) displacement, and F) reorientation. G-L) *mtnr1ab* cohort, with G) probability of response, H) response latency, I) maximum tail-bend amplitude, J) duration, K) displacement, and L) reorientation. M-R) *mtnr1al* cohort, with M) probability of response, N) response latency, O) maximum tail-bend amplitude, P) duration, Q) displacement, and R) reorientation.

**Figure S4.**
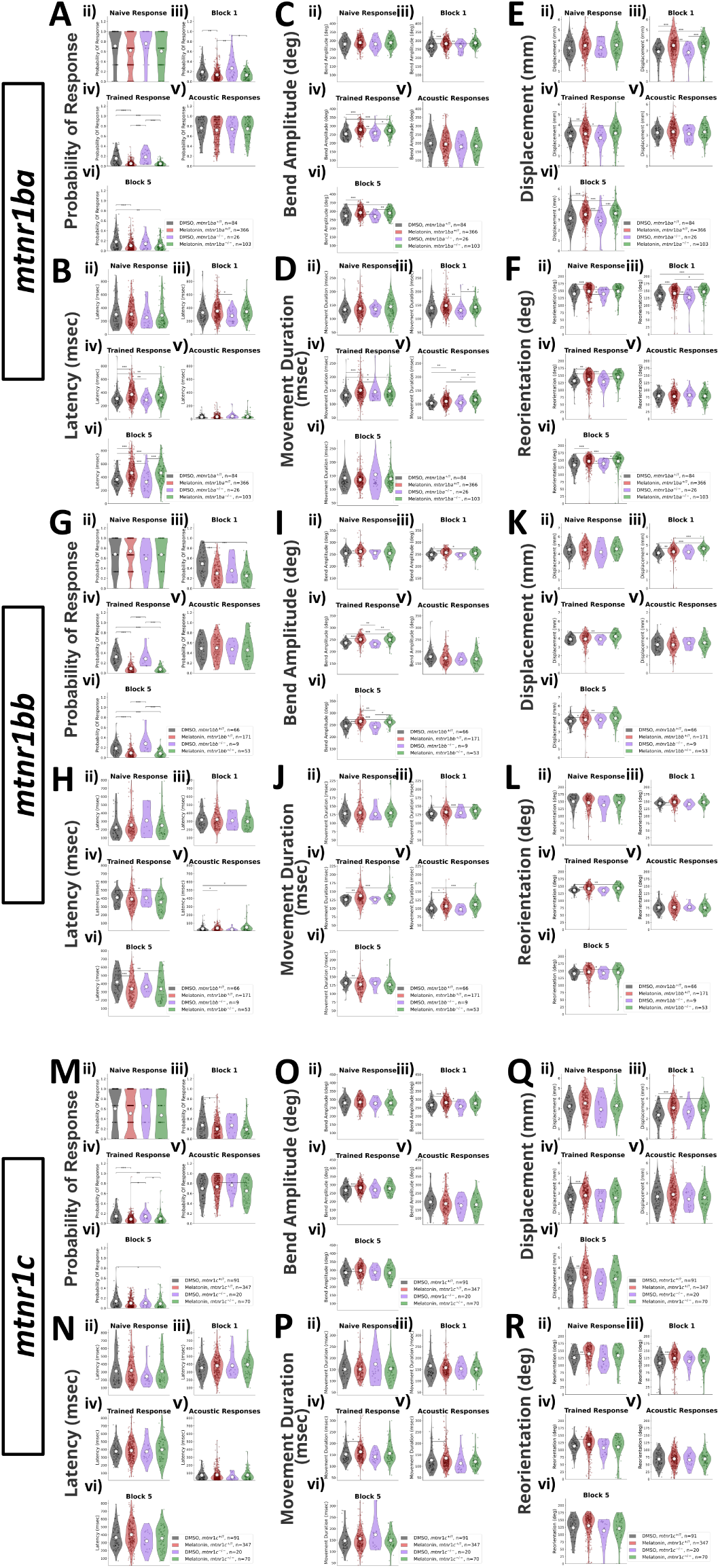
Quantification of epoch-level behavioural responses from Figure 4. Distributions of responsiveness for different epochs of the experiment ii)-v): ii) Naive Response (first 3 dark-flash stimuli), iii) Block 1 (remaining 57 stimuli of the first training block), iv) Trained Response (training blocks 2–4), and v) Acoustic Response (30 vibration stimuli). Each dot is the per-fish average of the epoch. Statistical significance was calculated using the Kruskal-Wallis test followed by Dunn’s test for post-hoc pairwise comparisons with Bonferroni correction, * = p < 0.05, ** = p < 0.01, *** = p < 0.001. A-F) *mtnr1ba* cohort, with A) probability of response, B) response latency, C) maximum tail-bend amplitude, D) duration, E) displacement, and F) reorientation. G-L) *mtnr1bb* cohort, with G) probability of response, H) response latency, I) maximum tail-bend amplitude, J) duration, K) displacement, and L) reorientation. M-R) *mtnr1c* cohort, with M) probability of response, N) response latency, O) maximum tail-bend amplitude, P) duration, Q) displacement, and R) reorientation.

**Figure S5.**
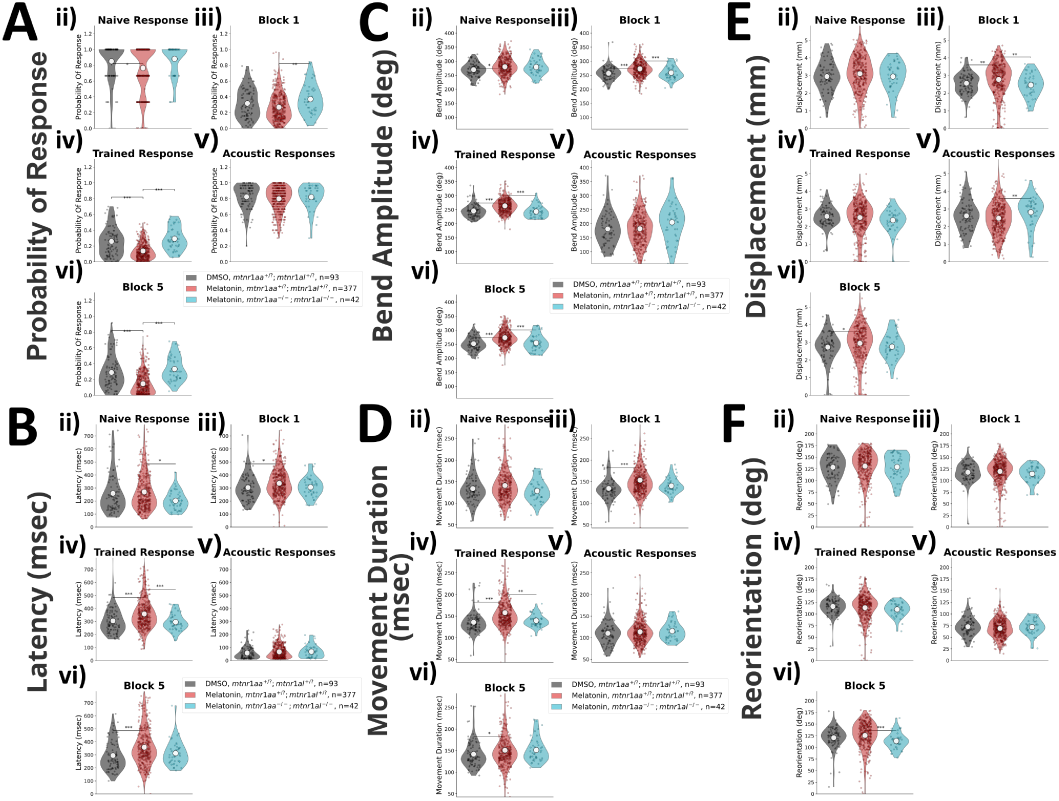
Quantification of epoch-level behavioural responses from Figure 5. Distributions of responsiveness for different epochs of the experiment ii)-v): ii) Naive Response (first 3 dark-flash stimuli), iii) Block 1 (remaining 57 stimuli of the first training block), iv) Trained Response (training blocks 2–4), and v) Acoustic Response (30 vibration stimuli). Each dot is the per-fish average of the epoch. Statistical significance was calculated using the Kruskal-Wallis test followed by Dunn’s test for post-hoc pairwise comparisons with Bonferroni correction, * = p < 0.05, ** = p < 0.01, *** = p < 0.001. A-F) *mtnr1aa;mtnr1al* double-mutant experiment, with A) probability of response, B) response latency, C) maximum tail-bend amplitude, D) duration, E) displacement, and F) reorientation.

## Notes

https://github.com/owenrandlett/2026_MtnRs_Hab

### Competing Interest Statement

The authors have declared no competing interest.

